# Robust Small Molecule-Aided Cardiac Reprogramming Systems Selective to Cardiac Fibroblasts

**DOI:** 10.1101/2023.09.20.558597

**Authors:** Yanmeng Tao, Yang Yang, Zhenghao Yang, Lipeng Wang, Shi-Qiang Wang, Yang Zhao

**Affiliations:** State Key Laboratory of Natural and Biomimetic Drugs, Ministry of Educational Key Laboratory of Cell Proliferation and Differentiation, Beijing Key Laboratory of Cardiometabolic Molecular Medicine, Peking-Tsinghua Center for Life Sciences, Institute of Molecular Medicine, College of Future Technology, Peking University, Beijing 100871, China; State Key Laboratory of Membrane Biology, College of Life Sciences, Peking University, Beijing 100871, China

**Keywords:** cardiac reprogramming, selectivity, robust, transcription factors, chemicals, mouse, human

## Abstract

Direct cardiac reprogramming to induce cardiomyocyte-like cells, e.g. by GMT (Gata4, Mef2c and Tbx5), is a promising route for regenerating damaged heart *in vivo* and disease modelling *in vitro*. Supplementation with additional factors and chemical agents can enhance efficiency but raises concerns regarding selectivity to cardiac fibroblasts and complicates delivery for in situ cardiac reprogramming. Here, we screened 2000 chemicals with known biological activities and found that a combination of 2C (SB431542 and Baricitinib) significantly enhances cardiac reprogramming by GMT. Without Gata4, MT (Mef2c and Tbx5) plus 2C could selectively reprogram cardiac fibroblasts with enhanced efficiency, kinetics, and cardiomyocyte function. Moreover, 2C significantly enhanced cardiac reprogramming in human cardiac fibroblasts. 2C synergistically enhances cardiac reprogramming by inhibiting Alk5, Tyk2 and downregulating Oas2, Oas3, Serpina3n and Tgfbi. 2C enables selective and robust cardiac reprogramming that can greatly facilitate disease modelling *in vitro* and advance clinical therapeutic heart regeneration *in vivo*.

## INTRODUCTION

Due to the limited regenerative capacity of the adult mammalian heart^1,2^, direct cardiac reprogramming has emerged as a promising therapeutic strategy to replenish injured heart tissues by converting scar (fibroblasts) into functional heart cells (cardiomyocytes)^3–18^. However, low efficiency in the induction of functional cardiomyocytes has hampered its broad application. To address this issue, several groups have developed various cocktails by adding more transcription factors to GMT, such as Hand2^6^, Hand2+Nkx2.5^19^, Akt1+Hand2^15^, PHF7^18^, Hand2+Nkx2.5^20^. Different from GMT in mouse cells, cardiac reprogramming in human cells requires additional reprogramming factors^4,8–10,12^, but with increasing numbers of required factors, the difficulty of *in vivo* delivery also increases.

Compounding the complexity of this issue, different combinations of GMT-based cardiac reprogramming factors have been used to successfully reprogram a broad range of fibroblasts, such as those derived from mouse embryos^19–22^, tail-tips^3,13,15,16,22–25^, raising the concern that direct delivery of GMT *in vivo* may lead to severe off-target reprogramming of cells in other organs into beating cardiomyocytes. Although supplementing additional transcription factors to GMT is a reasonable strategy to enhance cardiac reprogramming, the increased difficulty of *in vivo* delivery and the increased risks for reprogramming cells other than cardiac fibroblasts should be taken into consideration. A robust and cardiac fibroblast-specific reprogramming system to induce cardiomyocytes is thus required to advance the therapeutic application of this technology *in vivo*.

Cardiogenic genes (i.e., Gata4, Mef2c, Tbx20) expressed in cardiac fibroblasts reportedly contribute to heart development and repair^26^, suggesting that the cardiogenic signature of cardiac fibroblasts can be leveraged to identify a well-tailored combination of reprogramming factors that selectively reprograms cardiac fibroblasts, but not fibroblasts derived from other tissues or organs.

Small molecules have been widely used in the manipulation of cell fates or identities. For instance, they have been employed to induce chemically induced pluripotent stem cells (CiPSCs)^27–29^, chemically induced neurons (CiNs)^30,31^ extended pluripotent stem cells (EPSCs)^32^, chemically induced human pluripotent stem cells (hCiPS)^33^, and long-term maintenance of primary human hepatocytes *in vitro*^34^. Moreover, small molecules have been reported to enhance cardiac reprogramming efficiency. These include SB431542, an ALK5 inhibitor^20^; A83-01, a ALK4/5/7 inhibitor^22^; Y-27632, a ROCK1/2 inhibitor^22^; MM408, a Mll1 H3K4 methyltransferase inhibitor^35^; SB431542 combined with XAV939, a WNT inhibitor^4^; DAPT, a Notch inhibitor^17^; and IMAP, comprising IGF-1 (insulin-like growth factor-1), MM589 (a Mll1 inhibitor), A83–01 (a transforming growth factor-beta inhibitor), and PTC-209(a Bmi1 inhibitor)^36^.

In this study, we developed a high-efficient cardiac reprogramming cocktail, 2C+MT, that selectively reprogramed cardiac fibroblasts, but showed no such effects in fibroblasts from skin, tail-tip, brain, lung, or liver. These findings open possibilities to induce in situ direct reprogramming without harming irrelevant cell types.

## RESULTS

### The 2C combination boosts cardiac reprogramming

To identify chemicals that can enhance cardiac reprogramming, we screened neonatal skin fibroblasts^37^ (NSF) isolated from transgenic mice expressing mCherry driven by the cardiomyocyte-specific promoter, ɑMHC^3,38^. In total, ∼2000 compounds from both commercially available and proprietary in-house libraries were tested for their effects in NSF cells infected with GMT (Gata4, Mef2c, and Tbx5)-expressing lentivirus. The potentially induced cardiomyocyte-like cells (iCMs) were detected by activation of ɑMHC-mCherry at two weeks post treatment (Figure 1A). Reprogramming efficiency was quantified by counting mCherry-positive cells or immunostaining instead of flow cytometric analysis, since in situ counting cell numbers can significantly get rid of false positive cells by checking cardiomyocyte-like morphology (rod-shaped with increased size than Ctrl or unconverted fibroblasts or even with well-organized sarcomere), and by avoiding the bias of trypsin that selectively dissociates fibroblasts than iCMs before flow cytometric sorting analysis (Movie S1). This screen identified two compounds that appeared to synergistically enhance cardiac reprograming. In the DMSO-treated control group, approximate quantification by fluorescence microscopy revealed an average of 5 ɑMHC-mCherry+ cells in each field of view, while an average of 13.7 ɑMHC-mCherry+ cells were detected in the C1 (SB431542) group, and an average of 87.3 induced cells per field of view were observed in the C2 (Baricitinib) group. By contrast, 178.3 ɑMHC-mCherry+ cells per field of view were observed in the 2C (SB431542 and Baricitinib) combination group (Figure 1B-C). In wild-type NSF (Figure 1D), we performed immunofluorescence staining for ɑ-actinin (depicted in Figure 1E and quantified in 1F).

**Figure 1.**
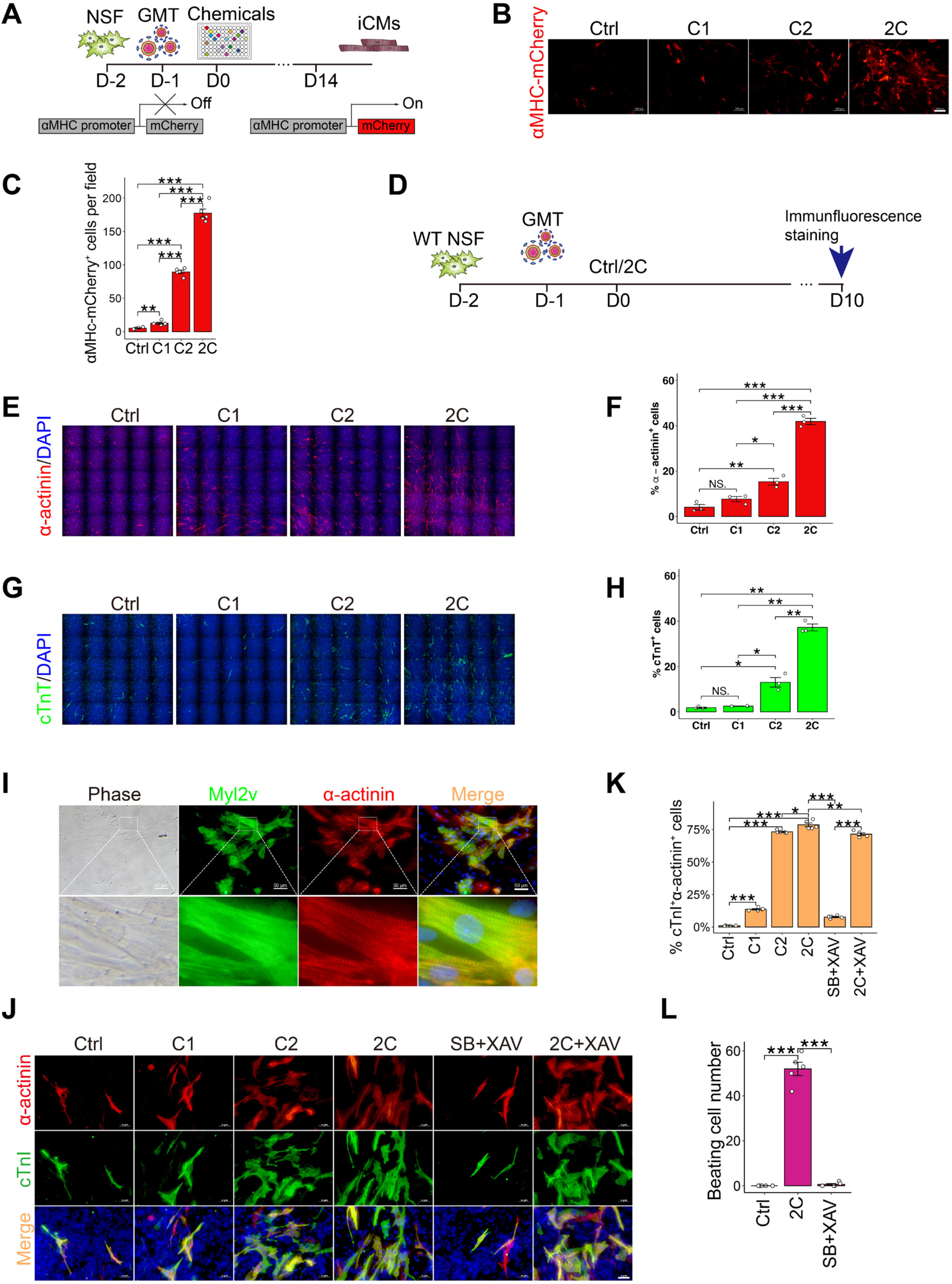
The 2C combination boosts cardiac reprogramming

Our findings indicate that 2C+GMT induced 41.9% ɑ-actinin+ cells, whereas the NSF control, DMSO control, C1, and C2 treatments resulted in substantially lower proportions of ɑ-actinin+ cells at 0%, 4.1%, 7.6%, and 15.3%, respectively. In Figure 1G and 1H, 2C+GMT induced a 37.2% increase in cTnT+ cells, whereas the NSF control, DMSO control, C1, and C2 treatments resulted in lower percentages of cTnT+ cells at 0%, 1.8%, 2.5%, and 13%, respectively. Furthermore, the large majority of iCMs induced by G2C+MT co-expressed Myl2v (myosin light chain 2) and ɑ-actinin and had a well-defined sarcomere structure (Figure 1I).

Further optimization showed that 2 µM concentrations of C1 and C2 were the most effective dosage for inducing the highest proportion of cTnT+ cells (Figure S1A-D), while the highest efficiency was obtained through continuous treatment during induction, indicated by the high percentage of ɑ-actinin+ (sarcomeric alpha actinin) cells and the number of spontaneous beating cells (a reliable readout of iCMs functionality) (Figure S1E). Based on our evidence of consistent co-expression of cTnI (cardiac troponin I), cTnT, and ɑ-actinin in iCMs (Figure S2A-B), we used these markers interchangeably throughout our study due to the low availability of each antibody.

It was previously reported that the SB431542+XAV939 (SB+XAV) combination could significantly increase cardiac reprogramming efficiency^4^. In comparison with 2C, SB+XAV showed moderate improvement on the percentage of cTnI+ and ɑ-actinin+ cells (Figure 1J, quantified in 1K). Regarding the number of spontaneous beating cells, 2C was much more efficient (Figure 1L, supplementary Movie S2-S4). Taken together, these results showed that the 2C cocktail could boost the efficiency of mouse cardiomyocyte reprogramming over that of vehicle control.

### 2C+MT selectively reprograms cardiac fibroblasts

Besides the above-mentioned great improvement of cardiac reprogramming in skin fibroblasts, we also tested the effect of 2C in cardiac fibroblasts. Adult mouse cardiac fibroblasts and neonatal mouse cardiac fibroblasts were isolated according to a previously described protocol^39^ and confirmed by immunofluorescence staining of cTnT and IZl-actinin to exclude any contamination with cardiomyocytes (Figure S3). Consistent with previous report^26^, our results showed that several cardiogenic genes (i.e., Gata4, Mef2c, Tbx20, Hand2, Nkx2-5) expressed in adult mouse cardiac fibroblasts (ACF) compared to adult mouse skin fibroblasts (ASF) (Figure 2A). Inspired by this result, 2C was tested with each individual transcription factor and each combination of two factors. Using the cTnT marker, we found that 2C could also enhance cardiac reprogramming in cells infected with only MT (Mef2c and Tbx5)-expressing lentivirus, but not other combinations (Supplemental Figure S4A, quantified in S4B). The combination of MT with 2C resulted in the induction of 15% IZl-actinin+ cells and 13.4% cTnT+ cells (shown in Figure 2B and quantified in Figure 2C and 2D). In contrast, treatments with GMT alone yielded only 0.6% IZl-actinin+ cells and 0.4% cTnT+ cells. To further fine-tune the stoichiometry of Mef2c (M) and Tbx5 (T), we expressed the M and T transcription factors in distinct ratios (Figure S5A-B) using a 2A “self-cleaving” peptide system. 2A peptides are short (18-22 amino acids) viral oligopeptides that mediate cleavage of polypeptides during translation in eukaryotic cells^40–42^. We found that the bi-cistronic M-T2A-T combination resulted in higher reprogramming efficiency than T-P2A-M, M-P2A-T or M-E2A-T, based on the increased percentage of ɑ-actinin+ cells and the greater number of spontaneous beating cells (Figure S5C-E). Further comparison of the 2C+(M-T2A-T) treatment with the tri-cistronic MGT construct^24^, in various combinations with 2C and/or SB+XAV, showed that the 2C+M-T2A-T combination induced significantly more spontaneous beating cells than any other combination, including 2C+MGT (Figure 2E, Movie S5-S6).

**Figure 2.**
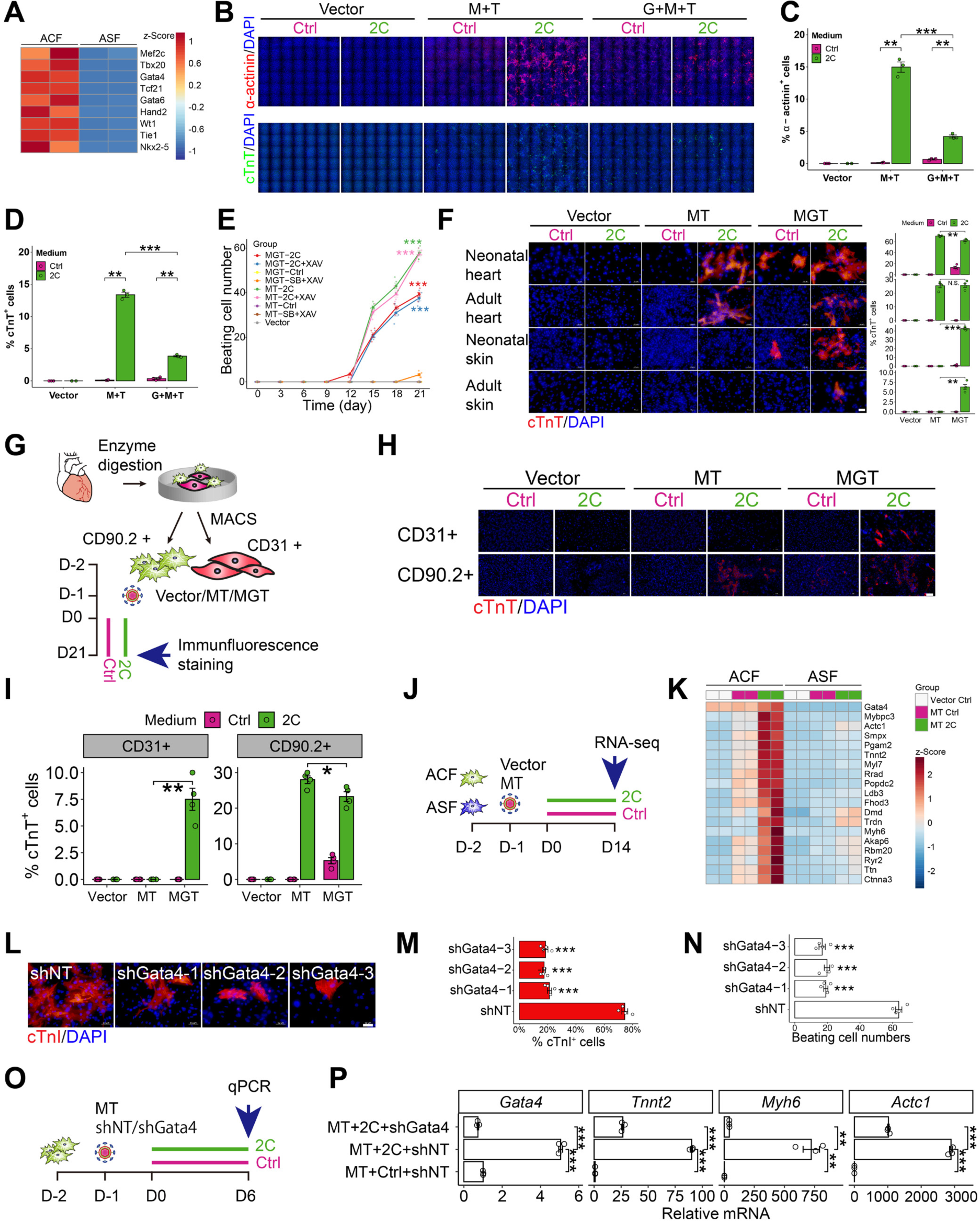
2C+MT selectively reprograms cardiac fibroblasts

Testing a range of cell types showed that 2C+MT (2C+M-T2A-T) could selectively reprogram cTnT+ iCMs from mouse neonatal cardiac fibroblasts and adult cardiac fibroblasts at 70.5% and 25.5% respectively (Figure 2F). However, either negligible or no reprogramming effects of 2C+MT were observed in mouse neonatal skin fibroblasts or adult skin fibroblasts. By contrast, 2C+MGT treatment led to efficient reprogramming in all these fibroblast types. We tested various fibroblasts derived from different tissues/organs (skin, heart, brain, lung, liver) with consistent results (Figure S6). To further determine if 2C+MT selectively reprogram cardiac fibroblasts, we sorted CD90.2+ cardiac fibroblasts and CD31+ endothelial cells by MACS (magnetic-activated cell sorting) and performed iCMs reprogramming (Fig2G). Our result showed that 2C+MT efficiently induced iCMs from CD90.2+ cardiac fibroblasts, while no cTnT+ iCMs could be detected in CD31+ endothelial cells (Figure 2H, quantified in 2I). These results indicated that 2C+MGT might be widely applicable for *in vitro* cell reprogramming in different fibroblasts, especially skin fibroblasts, which are easily accessible. However, the markedly stronger selectivity of 2C+MT suggested its potential *in vivo* therapeutic application for expanding beating cardiomyocytes while avoiding fibroblasts conversion in other tissues, such as skin.

To further validate the selectivity of 2C+MT in cardiac fibroblast, we performed RNA-seq in ACF and ASF (Figure 2J). 2C+MT treatment led to up-regulation of a panel of cardiac genes (i.e., Gata4, Actc1, Tnnt2, Myh6, Ryr2, and Ttn) in cardiac fibroblasts, while no significant up-regulation of these genes was detected in skin fibroblasts (Figure 2K). To validate the hypothesis that 2C+MT enables selective cardiac reprogramming in cardiac fibroblasts dependent on the expression of endogenous cardiogenic genes, we induced Gata4 knockdown in neonatal mouse cardiac fibroblasts by shRNA, which resulted in significantly fewer cTnI+ (Figure 2L, quantified in 2M) and spontaneous beating cells (Figure 2N) induced by 2C+MT compared to that in the non-target shRNA controls (shNT). Moreover, we performed qPCR 6 days post induction to determine the expression of cardiac genes (Figure 2O). 2C+MT significantly up-regulated the expression of Gata4, Tnnt2, Myh6, and Actc1, all of which were down-regulated by the knockdown of Gata4 (Figure 2P). These results indicated that endogenous cardiogenic genes (including but not limited to Gata4) expression in cardiac fibroblasts can be further enhanced/activated by 2C to enable selective cardiac reprogramming.

To determine if both C1 and C2 are essential to substitute Gata4 in the GMT treatment, we treated mouse neonatal cardiac fibroblasts infected with MT-expressing lentivirus with C1, C2, 2C, SB+XAV, 2C+XAV (2C+XAV939), or DMSO (Ctrl). Quantification of cTnT+ cells and spontaneous beating cells revealed that MT in combination with either C1, C2, or SB+XAV could induce cTnT+ cells at low efficiency (Figure S7A-B), but spontaneous beating iCMs could only be detected in the presence of 2C, and that XAV did not further enhance the effects of 2C (Figure S7C). These results thus indicated that 2C synergistically boosts the efficiency and kinetics of MT-mediated cardiac reprogramming, selectively in cardiac fibroblasts, and is essential for inducing spontaneous beating cells.

### 2C+MT-reprogrammed iCMs functionally and transcriptomically resemble adult cardiomyocytes

Morphological observations confirmed that sarcomere structure was well-organized in iCMs induced by 2C+MT (Figure 3A,Movie S7), while 2C+MT or 2C+GMT induced iCMs with spontaneous calcium transients (Figure 3B, Movie S8) and action potential (Figure 3C). To better understand how 2C affects cardiac fibroblasts reprogramming, we used RNA-seq for transcriptomic profiling of iCMs induced by 2C or other combinations. In total, 1498 genes were differentially up- regulated in 2C treatments while 2085 genes were differentially down-regulated at 5 weeks post induction (Figure 3D). Transcriptomic profiling showed that 2C treatment led to significantly higher cardiomyocyte-specific gene expression than that in control groups (i.e., no 2C or SB+XAV) (Figure 3E), while fibroblast-related genes were significantly down- regulated compared with their expression in SB+XAV treatments. Subsequent qPCR analysis of cardiomyocyte identity and functional markers also confirmed that iCMs derived from mouse neonatal cardiac fibroblasts expressing MT, GMT, or the empty vector control, exposure to 2C resulted in significant up-regulation of the cardiomyocyte-specific sarcomere genes (Myh6, Tnnt2 and Actc1) and the functional markers (Ryr2, Nppa and Gja1) (Figure 3F).

**Figure 3.**
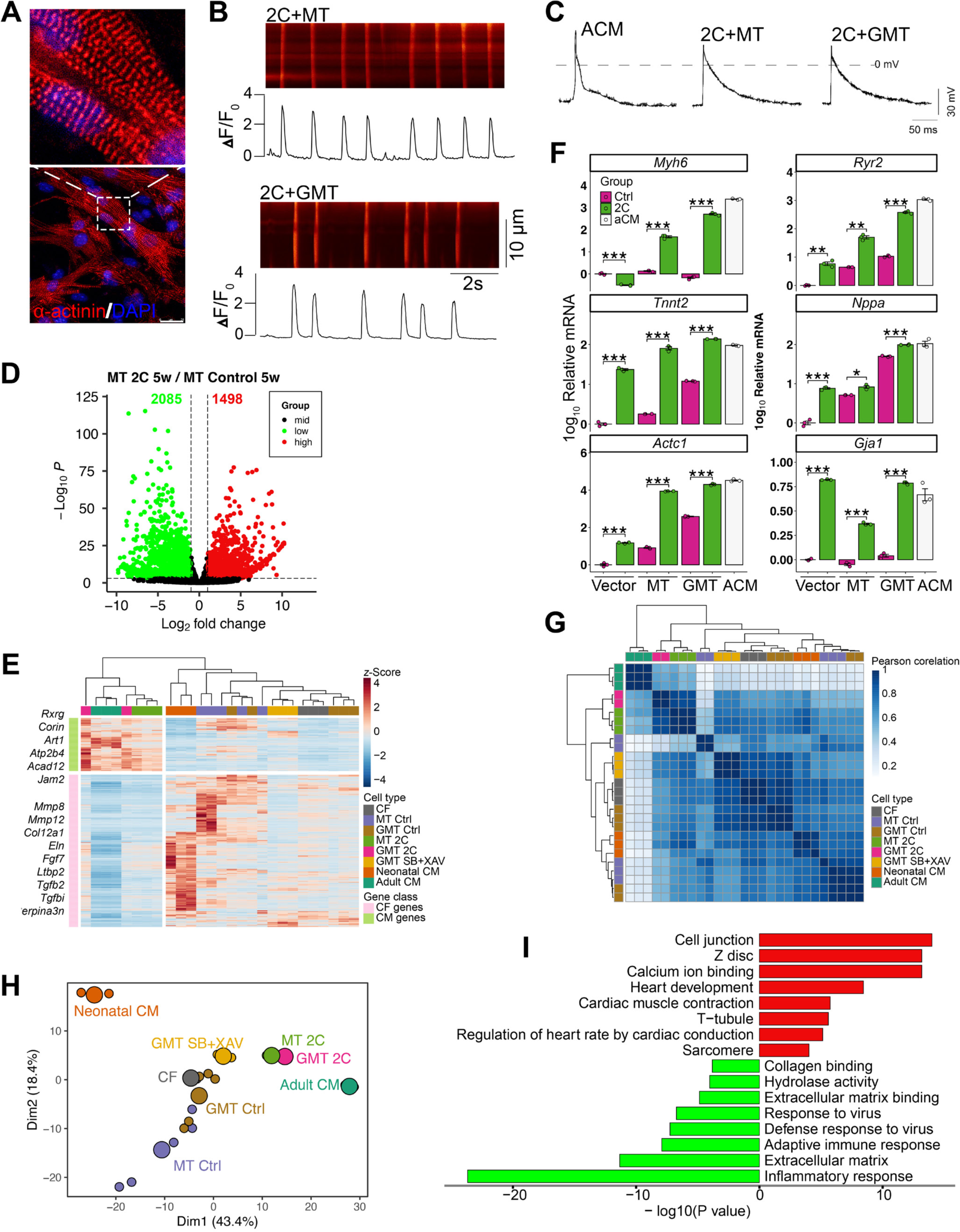
2C+MT-reprogrammed iCMs functionally and transcriptomically resemble adult cardiomyocytes

According to the Pearson corelation analysis (Figure 3G) and principal component analysis (PCA) (Figure 3H), 2C promoted fibroblast conversion toward an adult cardiomyocyte gene program compared with GMT/MT alone or GMT/MT+SB431542+XAV939. Gene ontology (GO) term enrichment analyses suggested that the up-regulated genes were associated with cell junction, Z-disc, calcium ion binding, heart development, cardiac muscle contraction, T-tubule and sarcomere, while the downregulated genes were associated with inflammatory response, extracellular matrix, and response to virus (Figure 3I). Furthermore, 2C enhanced cardiac reprogramming progressively (Figure S8A-E). These cumulative results implied that 2C treatment could enhance functionality in reprogrammed iCMs, most likely through inducing an adult cardiomyocyte signature, resulting in calcium transients and action potential.

### 2C enhance cardiac reprogramming in human cardiac fibroblasts

Direct cardiac reprogramming in human cells is substantially more complicated than in mouse fibroblasts, requiring more reprogramming factors and longer induction time, while resulting in lower reprogramming efficiency and less functional state^4,8–10,12^. Since 2C could increase the efficiency, kinetics, and selectivity of cardiac reprogramming in mouse cells while reducing the number of requisite transgenes and improving functionality in iCMs, we then evaluated the effects of 2C in reprogramming human cardiac fibroblasts. To this end, we transduced human cardiac fibroblasts with GATA4, MEF2C, TBX5, MESP1 and MYOCD (5F), as described by Wada and colleagues^10^, and treated the cells with solvent control (Ctrl) or 2C. Human cardiac fibroblasts we used were confirmed by the expression of Periostin and without expression of ɑ- actinin, cTnT or cMHC (Figure S9). Immunofluorescence staining for ɑ-actinin (depicted in Figure 4A and quantified in 4B) revealed that 2C+5F induced 26.6% of ɑ-actinin+ cells, whereas the DMSO control resulted in only 7.4% ɑ-actinin+ cells 3 weeks post-induction. Similarly, immunofluorescence staining for cTnT (depicted in Figure 4A and quantified in 4C) demonstrated that 2C+5F induced 7.7% cTnT+ cells, whereas the DMSO control resulted in only 2.4% cTnT+ cells. Additionally, qPCR analysis indicated a significant upregulation of cardiac genes (i.e., MYH6, TNNT2, ACTN2, and ACTC1) in the 2C+5F-induced cardiomyocytes (Figure 4D). These findings collectively demonstrate the efficacy of 2C in enhancing the reprogramming of human cardiac fibroblasts. Furthermore, when employing 2C+4F (comprising MEF2C,

**Figure 4.**
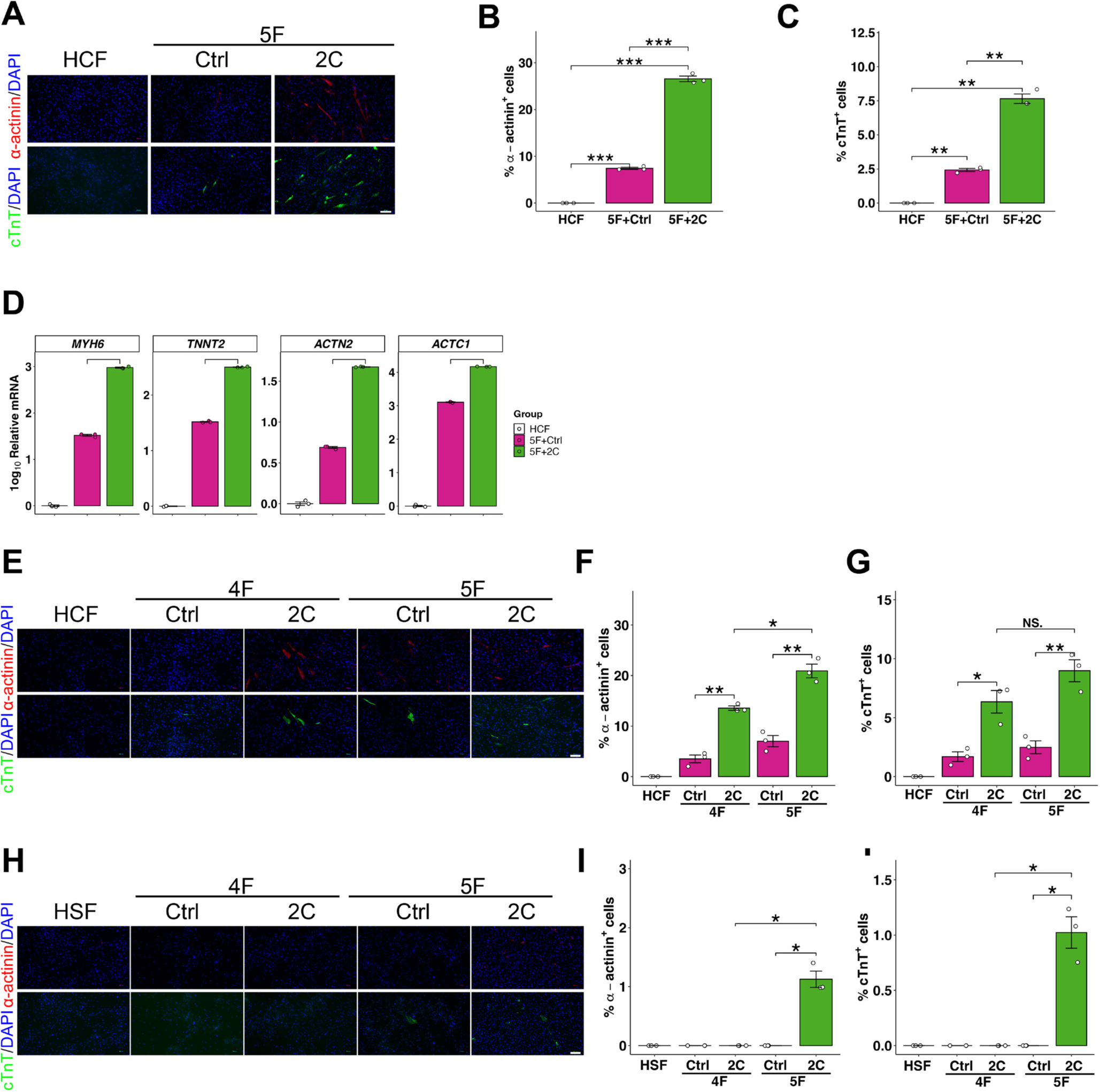
2C enhance cardiac reprogramming in human cardiac fibroblasts

TBX5, MESP1, and MYOCD), we observed an induction of 13.6% ɑ-actinin+ cells (depicted in Figure 4E and quantified in 4F) and 6.3% cTnT+ cells (depicted in Figure 4E and quantified in 4G) in human cardiac fibroblasts (HCF). In contrast, negligible or no reprogramming effects (Figure 4H-J) of 2C+4F were observed in human skin fibroblasts (HSF). Taken together, our results indicate that 2C+5F significantly enhance cardiac reprogramming in both human cardiac and skin fibroblasts, whereas 2C+4F selectively reprogram human cardiac fibroblasts.

### 2C increases the proportion of iCMs and inhibits cell proliferation

Considering our above data showing that 2C can increase the efficiency and functionality of cardiac reprogramming, we then investigated whether and how the effects of 2C might be related to cell proliferation by testing three models (Figure 5A). First, we hypothesized that 2C enhanced the proliferation of converted iCMs. Second, we speculated that 2C could promote proliferation in the total cell population. Our third hypothesis was that 2C directly increased the conversion ratio. To test these hypotheses, we conducted immunofluorescence staining to simultaneously quantify proliferating cells (indicated by Cyclin D1+ cells), iCMs (indicated by LJ-actinin+ cells), and total cell numbers (DAPI+ cells) at various time points over 16 days (Figure 5B). These assays showed that 2C enhanced cardiac reprogramming efficiency in a time-dependent manner by increasing the absolute number of LJ-actinin+ iCMs (Figure 5C-D), inhibiting cell proliferation (Figure 5E), decreasing the absolute number of total cells (Figure 5F) and thus resulting in the more higher ratio of iCMs to total cells (Figure 5G). No Cyclin D1 and LJ-actinin double positive cells could be detected throughout the induction (Figure 5H).

**Figure 5.**
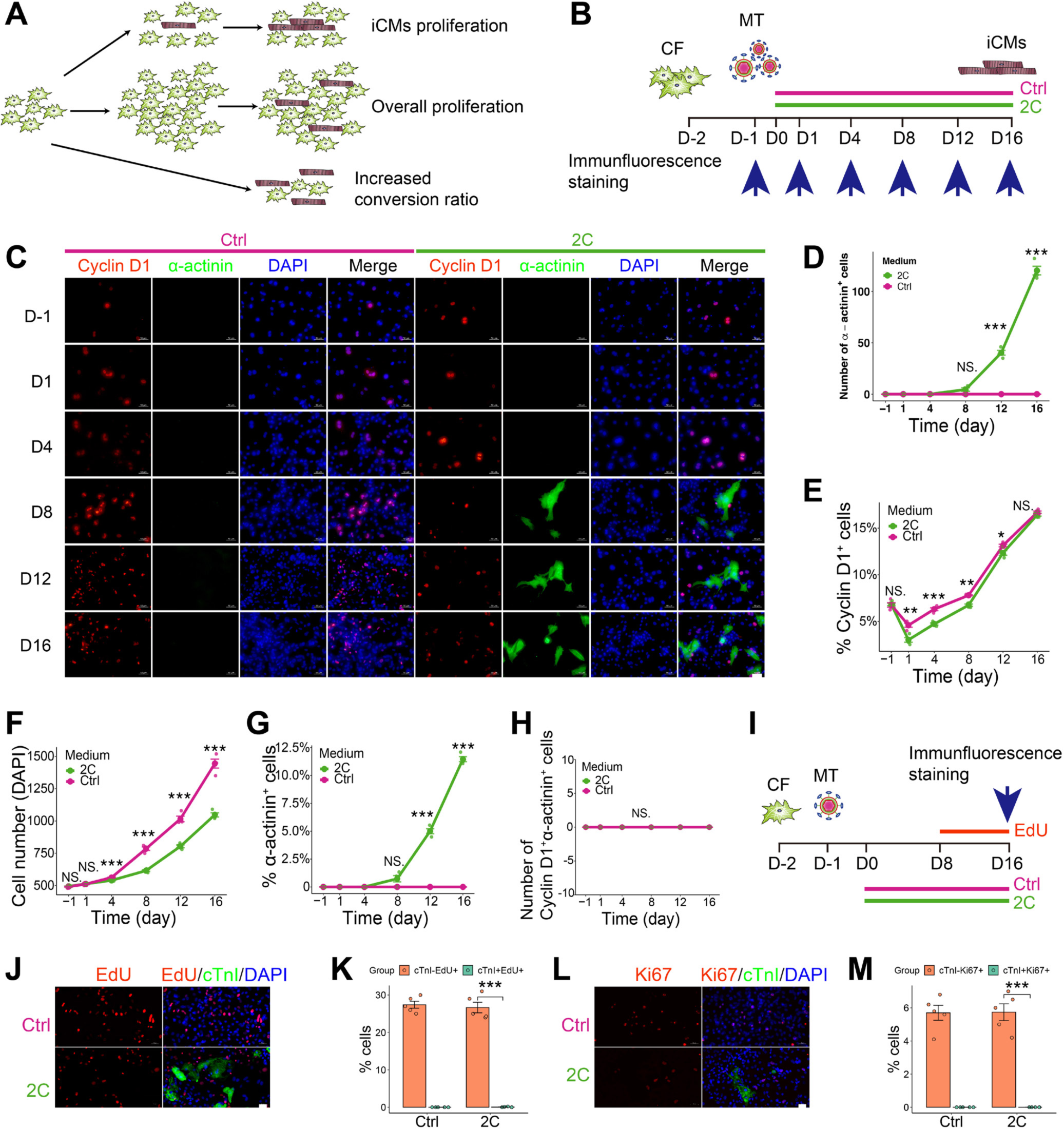
2C increases the proportion of iCMs and inhibits cell proliferation

We also performed EdU incorporation (Figure 5I-K) and Ki67 staining (Figure 5L-M) to further confirm that 2C+MT induced iCMs without inducing iCMs proliferation. These cumulative data indicated that 2C enhances cardiac reprogramming by increasing the absolute number of iCMs while inhibiting cell proliferation and thus increasing the conversion ratio.

### 2C+MT-reprogrammed iCMs maintain a stable cardiomyocyte fate

In addition to optimizing the 2C treatment duration (Figure S1E), we also asked whether the iCMs retained their cardiomyocyte-like phenotype in the absence of 2C. To answer this question, mouse cardiac fibroblasts were infected with MT-expressing lentivirus and treated with 2C for 3 weeks, after which they were cultured for another 2 weeks with or without 2C (Figure 6A). Immunofluorescent staining revealed that the reprogrammed iCMs indeed maintained their cardiomyocyte-like fate, indicated by the stable number of cTnI+ cells detected in cultures without 2C (Figure 6B-C). By contrast, the number of total cells rapidly increased in the absence of 2C (Figure 6D), resulting in a decreased proportion of cTnI+ cells (Figure 6E). These data indicated that the fibroblasts which did not undergo reprogramming began to rapidly proliferate following withdrawal of 2C treatment. Further, the iCMs induced by 2C+MT were stably reprogrammed and did not require 2C to maintain a persistent cardiomyocyte phenotype.

**Figure 6.**
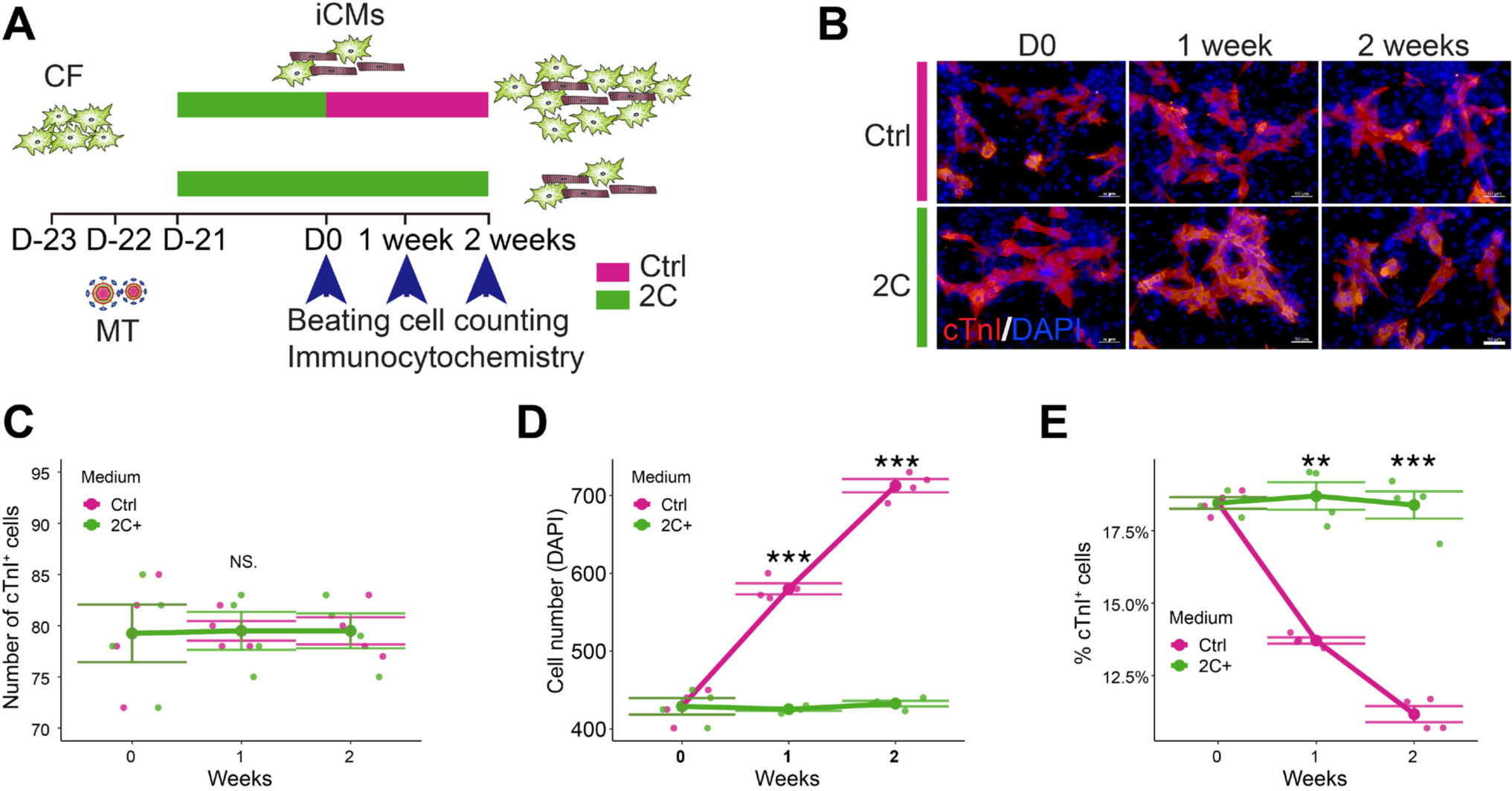
2C+MT induced iCMs maintain a stable cardiomyocyte fate

### C2 synergizes cardiomyocyte induction via suppression of C1-activated molecular barriers

Alk5 (Transforming Growth Factor Beta Receptor 1, also known as Tgfbr1) stands as a well-established target for C1^43,44^, while C2 has been associated with targeting Jak1, Jak2, Jak3, and Tyk2^45^. To discern the impact of 2C on the reprogramming process through these canonical targets, we conducted tests involving various chemicals known to share these regulatory targets with C1 and C2, respectively. Our findings revealed that all the chemicals tested, with the same targets as C1, significantly enhanced cardiac reprogramming efficiency, as indicated by the proportion of cTnT+ cells (Figure S10A). These results align with previous reports emphasizing the role of inhibiting the TGF-beta signaling pathway in cardiac reprogramming^4,20,22^.

Furthermore, we aimed to mimic the effects of C1 by conducting knockdown experiments targeting the receptors Alk4 and Alk5. Alk4 knockdown did not exhibit any enhancement of cardiac reprogramming. However, Alk5 knockdown yielded a substantial increase in the percentage of IZl-actinin+ cells, indicating a clear enhancement of cardiac reprogramming (Figure 7A-B). Notably, under Alk5 knockdown conditions, no spontaneous beating cells were detected (Figure 7C), suggesting that C1 may promote cardiac reprogramming through additional targets. Conversely, overexpression of Alk5 or Tgfb1 (Transforming Growth Factor Beta 1) resulted in significantly lower reprogramming efficiency, evidenced by a reduced proportion of cTnI+ cells and a decrease in the number of beating cells (Figure 7D-F). These observations imply that the inhibition of Alk5 is crucial for cardiac reprogramming but does not entirely account for the enhanced effects elicited by C1.

**Figure 7.**
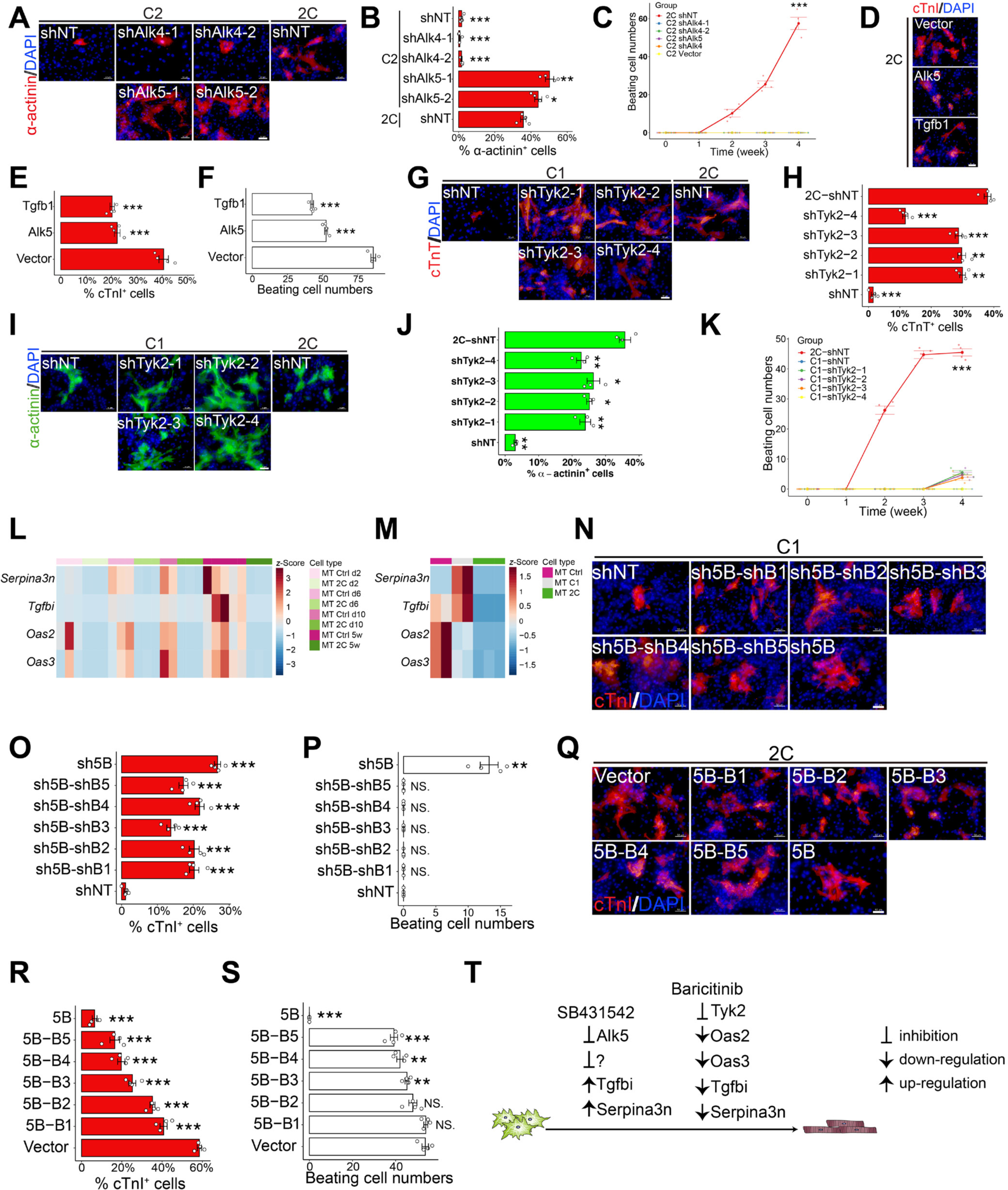
C2 synergizes cardiomyocyte induction via suppression of C1-activated molecular barriers

However, an investigation into the role of C2, coupled with JAK inhibitors, demonstrated that treatment with 19 out of 56 Jak inhibitors led to an increased percentage of cTnT+ cells, with 9 of them also significantly increasing the number of beating cells (Figure S10B-C). To genetically validate whether C2 enhances cardiac reprogramming through the inhibition of Jak1, Jak2, Jak3, or Tyk2, we conducted knockdown experiments using two independent shRNAs for each gene.

Knockdown of Jak1, Jak2, and Jak3 did not exert significant effects on the percentage of IZl-actinin+ cells, cTnT+ cells, or the number of spontaneous beating cells (Figure S11A-E). In contrast, Tyk2 knockdown resulted in an increased percentage of cTnT+ cells (Figure 7G-J). Notably, even after an extended induction period spontaneous beating cells were barely detectable (Figure 7K). Conversely, Tyk2 overexpression did not negate the effects of C2, even when used at various concentrations of C2 (Figure S11F-H).

Stat3 serves as a primary downstream target of Jak signaling, activated via phosphorylation of tyrosine residue at 705 (Y705)^43–47^ or serine residue at 727 (S727)^48–52^. To examine the role of Stat3, we introduced mutations in the Y705 and S727 residues to simulate the phosphorylated and dephosphorylated states of Stat3. Surprisingly, the percentage of IZl- actinin+ cells and the number of beating cells remained unaffected in both WT and mutated Stat3 (Figure S11I).

Subsequent RNA-seq analysis conducted on 2C+MT, C1+MT, or MT control fibroblasts over a five-week time course identified four genes - Oas2, Oas3, Serpina3n, and Tgfbi (Transforming Growth Factor Beta Induced) - that were significantly downregulated in the presence of 2C throughout the entire experimental period (Figure 7L). Intriguingly, Serpina3n and Tgfbi were upregulated in C1+MT cells but downregulated by 2C (Figure 7M). Immunofluorescence staining further demonstrated that the combined knockdown of these five genes, referred to as the “5B” set (Tyk2, Oas2, Oas3, Serpina3n, and Tgfbi), significantly increased the percentage of cTnI+ cells and the number of beating cells, underscoring their role as inhibitors of cardiac reprogramming (Figure 7N-P). In contrast, the overexpression of the 5B gene set effectively countered the effects of 2C+MT, resulting in a reduced percentage of cTnI+ cells and beating cells (Figure 7R-S). Moreover, in line with these findings in mouse cells, TGFBI, SERPINA3, and OAS3 were also significantly downregulated by 2C, while OAS2 did not exhibit significant downregulation (Figure S12). These data suggest that 2C may function similarly, although not identically, in human cardiac fibroblast (hiCM) reprogramming. Collectively, these results indicate that C1 enhances cardiac reprogramming through the inhibition of the TGF-beta signal pathway, consistent with previous reports^4,20,22,53^. C1 also exhibits some adverse effects that can inhibit cardiac reprogramming, but these effects are mitigated by C2, explaining their synergistic effects (Figure 1B-H). These cumulative findings demonstrate that the components of 2C function synergistically to enhance cardiac reprogramming by modulating multiple downstream effectors (Figure 7T).

## DISCUSSION

In this study, we found that 2C significantly enhances the efficiency of GMT-mediated reprogramming. Testing each individual transcription factor and each combination of two factors revealed that 2C+MT can selectively reprogram cardiac fibroblasts with significantly higher efficiency and kinetics than GMT. 2C+4F (MEF2C, TBX5, MESP1, and MYOCD) also selectively convert human cardiac fibroblasts into iCMs. Several cardiogenic genes, such as Gata4, are known to be expressed in cardiac fibroblasts^26^, potentially accounting for the high selectivity of this cocktail. Moreover, 2C further upregulates endogenous Gata4 expression in cardiac fibroblasts, but does not induce any detectable Gata4 expression in fibroblasts from other organs. In addition to 2C, Bmi1 knock-down^16^ or PHF7 over-expression^18^ can both complement the loss of Gata4 in reprogramming mouse cardiac fibroblasts, although further investigation is required to determine whether those factor combinations are also selective to cardiac fibroblasts. More recently, Ascl1+Mef2c was also reported to induce iCMs from cardiac fibroblasts, highlighting the cross-lineage potential of Ascl1, but shows poor selectivity towards cardiac fibroblasts, since MEF can also be reprogrammed^54^.

Our findings strengthen the basis for *in vivo* study or applications of direct cardiac reprogramming. Increased selectivity with a reduced number of requisite genetic factors may also facilitate therapeutic application of in situ direct cardiomyocyte induction from cardiac fibroblasts. Additionally, although evaluating the effects of 2C *in vivo* falls beyond the scope of the current study, we found that 2C significantly down-regulates inflammatory signalling activity, which is activated in damaged heart tissue^55,56^ These findings further support that 2C may be potentially applied in cardiac reprogramming *in vivo* to overcome inflammation-associated molecular barriers, aligning well with another study that showed cardiac reprogramming efficiency is enhanced via ZNF281-mediated suppression of inflammation^57^.

Although we found that 2C is required for the highest efficiency reprogramming in the first 3 weeks of cardiomyocyte induction, (Figure S1E), iCMs induced by 2C+MT are stably reprogrammed and do not require 2C to maintain a persistent cardiomyocyte phenotype (Figure 6). Thus, the short-term use of 2C might decrease potential toxicity and off-target effects, which can be leveraged to develop a feasible and safe therapeutic approach to repair damaged heart tissue *in vivo*.

C1 and C2 show synergistic effects in promoting cardiac induction. C1, an Alk5 inhibitor^58^, has gained widespread recognition for its role in inducing various cell types, including induced pluripotent stem cells (iPSCs)^59^, induced neurons (iNs)^31^, cardiac progenitor cells^60^, cardiac Purkinje cells^61^, and iCMs^4,14,20,53,62^. Notably, it has been demonstrated that inhibiting pro-fibrotic signaling, a major barrier to cardiac lineage reprogramming, can substantially enhance the reprogramming process^22^. Although Alk5 inhibition is essential for cardiac reprogramming, this function cannot fully explain the enhancement of reprogramming caused by C1, which thus requires further investigation. C2 is a pan-Jak inhibitor, and consistent with our study, two Jak inhibitors (Erlotinib and Ruxolitinib) can also reportedly enhance cardiac reprogramming, as indicated by elevated expression of cardiomyocyte markers^13^. Similarly, ZNF281 has been shown to improve cardiac reprogramming by modulating inflammatory gene expression^57^. These reports, in conjunction with our findings in this study, highlight the inhibitory role of Jak-Stat signalling in cardiac reprogramming. However, only a few pan-Jak inhibitors that share a structural similarity with C2 can enhance cardiac fibroblast reprogramming to efficiently induce spontaneous beating iCMs, indicating that more complicated mechanisms likely underpin C2 enhancement of iCMs. Here, we found that C2 promotes cardiac reprogramming via suppression of C1-activated molecular barriers, such as Tgbfi and Serpina3n, which may account for their synergistic effects.

In summary, we established a robust cardiac reprogramming system to convert mouse or human cardiac fibroblasts into cardiomyocytes selectively and efficiently, opening new potential avenues for clinical translation of targeted reprogramming by optimizing reprogramming cocktails and reducing the number of genetic factors.

## Limitations of the study

While we have reported the effectiveness of the cardiac fibroblast-selective reprogramming factor combination, 2C+MT, further investigations are warranted to ascertain the selectivity and efficiency of 2C+MT in reprogramming cardiac fibroblasts into cardiomyocyte-like cells in mouse models of myocardium infarction *in vivo*. Additionally, due to limitations related to the availability of human cardiac fibroblasts, this study did not determine the minimal factor combination required for cardiac reprogramming in the presence of 2C. Ultimately, although we have identified several factors as downstream effectors of 2C, their contributions to cardiac reprogramming need to be fully elucidated in the future.

## EXPERIMENTAL PROCEDURES

### Plasmid construction

Coding sequences of Gata4, Mef2c, and Tbx5 were amplified by PCR from pMXs-MGT (addgene, Cat#111810, a gift from the Qian lab). The FU-tet-o-hOct4 (addgene, Cat#19778) was cut by EcoR I. Inserts and linearized vector were assembled using a pEASY-Uni Seamless Cloning and Assembly Kit (TransGen, Cat#CU101). tetO-hGATA4 (addgene, #46030), tetO-hMEF2C (addgene, #46031), and tetO-hTBX5 (addgene, #46032) were purchased from addgene. FU-tet- o-hMESP1 and t FU-tet-o-hMYOCD were constructed in this work. Detailed list of plasmids is available at **KEY RESOURCES TABLE.** For each vector used in this article, we validated proper gene expression at desired levels (overexpression or knockdown, data not shown here).

### Viral packaging and transduction

Lentiviral vector (15 μg) together with pMDLg/pRRE, RSV/Rev, and VSV-G (5 μg for each), were co-transfected into 293T cells with the Ca_3_(PO_4_)_2_-method in 10-cm dishes and incubated 12-16 hours. On the following day, the medium was changed. 2 days after transfection, viral supernatant was collected and filtered through 0.45 μm filters (Millipore, Cat#SLHP033RB) and then placed onto target fibroblasts supplemented with 8 μg/mL polybrene (Sigma-Aldrich, Cat#H9268-10G). Lentiviruses were used freshly or frozen at -80°C for future use.

### Animals and surgery

All animal experiments were performed according to the Animal Protection Guidelines of Peking University and all procedures conformed to the guidelines from the NIH Guide for the Care and Use of Laboratory Animals. Transgenic mice expressing mCherry driven by the cardiomyocyte-specific promoter, LJMHC (B6;D2-Tg(Myh6*-mCherry)2Mik/J, The Jackson Laboratory, #021577), were a gift from Dr. Kotlikoff^38^. MI was performed in 5-week-old male ICR mice of the B6;129S4 genetic background (Jackson Laboratory #008214) induced by permanent ligation of the left anterior descending coronary artery (LAD) with a 7-0 prolene suture as described previously^63^. Euthanasia was performed by deep (4%) isoflurane anesthesia followed by exsanguination, consistent with American Veterinary Medical Association guidelines.

### Primary cell cultures

Isolated neonatal (1.5-day-old CD-1 mice) hearts were minced into small pieces less than 1 mm^3^ in size and digested with collagenase type II solution. After 4-6 days, cells were separated by magnetic cell sorting (MACS) to obtain CD90.2+ neonatal fibroblasts according to a previously described protocol^39^. Adult mouse (5 weeks) cardiac fibroblasts and fibroblasts from neonatal mouse lung, neonatal mouse liver, neonatal mouse brain, adult mouse skin and adult mouse lung were isolated using a similar protocol without MACS. MICF (cardiac fibroblasts isolated from adult mouse after myocardial infarction) were isolated from infarct and border regions of adult heart, 4-5 days after surgery. All fibroblasts used were confirmed by immunofluorescence staining to exclude any contamination with cardiomyocytes. Neonatal mouse cardiomyocytes were isolated according to a previously described protocol^64^. Adult mouse cardiomyocytes were isolated according to a previously described protocol^65^. Human cardiac fibroblasts were purchased from Lonza (Cat#CC2904). Human skin fibroblasts ware purchased from ATCC (Cat#CRL-2522).

### iCMs reprogramming

P1 neonatal mouse cardiac fibroblasts were plated into a 24-well plate at a density of 1 x 10^5^cells per well in FB medium (10% FBS in DMEM supplemented with 1% penicillin-streptomycin). On the following day, fibroblasts were transduced with MT (FU-tet-o-MT, FUdeltaGW-rtTA, MOI = 10) lentivirus in transfection medium (10% FBS in DMEM supplemented with 8 μg/mL polybrene). On the following day, the medium was changed to 2C medium (DMEM/M199 (4:1) supplemented with 10% FBS,10% KnockOut Serum Replacement, 1% MEM Non-Essential Amino Acids, 1% GlutaMAX, 1% penicillin-streptomycin, 2 µg/mL Doxycycline hyclate, 2 µM SB431542, 2 µM Baricitinib). Three days after infection, reprogramming medium was supplemented with 1 µg/mL Puromycin and 200 µg/mL Geneticin. The medium was changed every 3-4 days. For human cells, P6-P8 cells were plated into a 24-well plate at a density of 2 x 10^5^cells per well for reprogramming.

### Chemical screening

Neonatal mouse skin fibroblasts (NSF) were isolated from transgenic mice expressing mCherry driven by the cardiomyocyte-specific promoter, αMHC. NSF were plated into a 24-well plate at a density of 1 x 10^5^ cells per well in FB medium. The following day, NSF were infected with GMT-expressing lentivirus (FU-tet-o-Gata4, FU-tet-o-Mef2c, FU-tet-o- Tbx5 and FUdeltaGW-rtTA, ∼100 µL unconcentrated virus, MOI = 10) in transfection medium. After 1 day, the medium was changed to reprogramming medium with Ctrl (DMSO) or 2 µM indicated chemicals. The medium was changed every 2-3 days. Two weeks after induction, ɑMHC-mCherry expression was quantified by fluorescence microscopy. In total, we screened ∼2,000 compounds from both commercially available and proprietary in-house libraries.

### Immunocytochemistry

After fixation with 4% paraformaldehyde (DingGuo, Cat#AR-0211) at room temperature for 30 min, cells were permeabilized with PBS-0.3% Triton X-100 (Sigma-Aldrich, Cat#T8787) and blocked with blocking buffer (PBS supplemented with 2.5% donkey serum (Jackson Immuno Research, Cat#017-000-121)) at 37℃ for 1 hour. Primary antibody (Cardiac Troponin T, Thermo Fisher, Cat#MA5-12960; Cardiac Troponin I, abcam, Cat#ab56357; Sarcomeric Alpha Actinin, abcam, Cat#ab9465; Cyclin D1, abcam, Cat#ab16663; Myosin Light Chain 2, abcam, Cat#ab48003; Ki67, abcam, Cat#ab15580; Periostin, abcam, Cat#ab14041; cMHC, abcam, Cat#ab207926) incubation with appropriate dilutions were performed at 4℃ overnight in blocking buffer. The following day, cells were washed with PBS three times and probed with secondary antibodies at 37℃ for 1 hour in blocking buffer. Cells were then washed with PBS three times and DNA was stained with DAPI solution (Sigma-Aldrich, Cat#D9542). For the quantification, 3-5 independent experiments were used for scanning of whole wells and analysed in Image J software, or 5 fields were randomly selected in a blinded manner and the indicated cells were counted manually in each experiment.

### qPCR

Total RNA was isolated with RNeasy Plus Mini Kit (QIAGEN, Cat#74134) and reverse-transcribed into cDNA using TransScript One-Step gDNA Removal and cDNA Synthesis SuperMix (TransGen, Cat#AT311). qRT-PCR was performed using the q225 Real-Time PCR system (Kubo Tech, Cat#q225) and ChamQ SYBR qPCR Master Mix (Vazyme,

Cat#Q321-02) according to the manufacturer’s instructions. The primer sequences used for qRT-PCR are provided in supplementary Table S1. mRNA levels were normalized by comparison to Gapdh mRNA.

### Calcium imaging

Calcium imaging was performed according to the standard protocol. Briefly, cells were loaded with 10 µM fluo4-AM (invitrogen, Cat#F14201) for 5 min at 37℃ in the dark. Line-scan images were performed on a confocal microscope (Carl Zeiss, Cat#LSM-710) and acquired at sampling rates of 3.78 ms/line and 0.03 µm/pixel.

### Electrophysiological recording

Whole-cell patch clamping was applied for action potential recording by an Axon 200B patch-clamp amplifier (Axon Instruments, USA). The action potentials were recorded in current clamp mode. Pipette resistances were 1.5–3 MΩ. The external solution contained (in mmol/L): 140 NaCl, 5.4 KCl, 1.8 CaCl_2_,1 MgCl_2_, 10 Glucose, 10 HEPES (pH 7.4 adjusted with NaOH). The pipette solution contained 120 KCl, 1 MgCl_2_, 3 MgATP, 10 HEPES,10 EGTA at pH 7.2 adjusted with KOH.

### RNA-Seq and Transcriptome Analysis

Total RNA was isolated using a RNeasy Plus Mini Kit (QIAGEN, Cat#74134). Total amounts and integrity of RNA were assessed using the RNA Nano 6000 Assay Kit of the Bioanalyzer 2100 system (Agilent Technologies, CA, USA). Total RNA was used as input material for RNA sample preparations. Differential expression analysis was performed using the DESeq2 R package^66^ (1.36.0) and visualized using the EnhancedVolcano R package (1.14.0). Genes with an adjusted P value < 10-4 and fold-change > 2 were assigned as differentially expressed. Gene ontolgy (GO) term enrichment analyses were performed using the DAVID 6.8 functional annotation tool^67^. Terms that had a P-value < 0.05 were defined as significantly enriched.

### Statistical Analysis

Values were presented as means ± SEM. The unpaired t-test was used to determine the significance of differences between two groups. Statistical analysis was performed using R 4.2.1. A value of P < 0.05 was considered statistically significant (*), a P value of < 0.01 was considered highly significant (**), a P value of < 0.001 was considered very highly significant (***), and a P value of > 0.05 was labelled as n.s (not significant). All data are representative of multiple independent experiments.

## Supporting information

Table-S1-List-of-primers

## ACKNOWLEDGMENTS

We thank Li Qian (University of North Carolina, Chapel Hill) for kindly providing the pMXs-MGT plasmid and detailed cardiac reprogramming protocol. We thank Yu Nie (Fuwai Hospital) for kindly providing the αMHC-mCherry mice. This work is supported by the National Key Research and Development Program of China (2018YFA0800504), the National Natural Science Foundation of China (31922020), and fundings provided by Plastech Pharmaceutical Technology Co., LTD.

## AUTHOR CONTRIBUTIONS

Yanmeng Tao designed and performed experiments, analysed data; Yang Yang assisted with some data collection about human cardiac reprogramming; Zhenghao Yang assisted with chemical screening; LiPeng Wang and Shiqiang Wang contribute to a part of the calcium imaging and all electrophysiological recording; Yang Zhao conceived this work and supervised this study. Yanmeng Tao and Yang Zhao wrote the manuscript.

---

**Figure S1.**
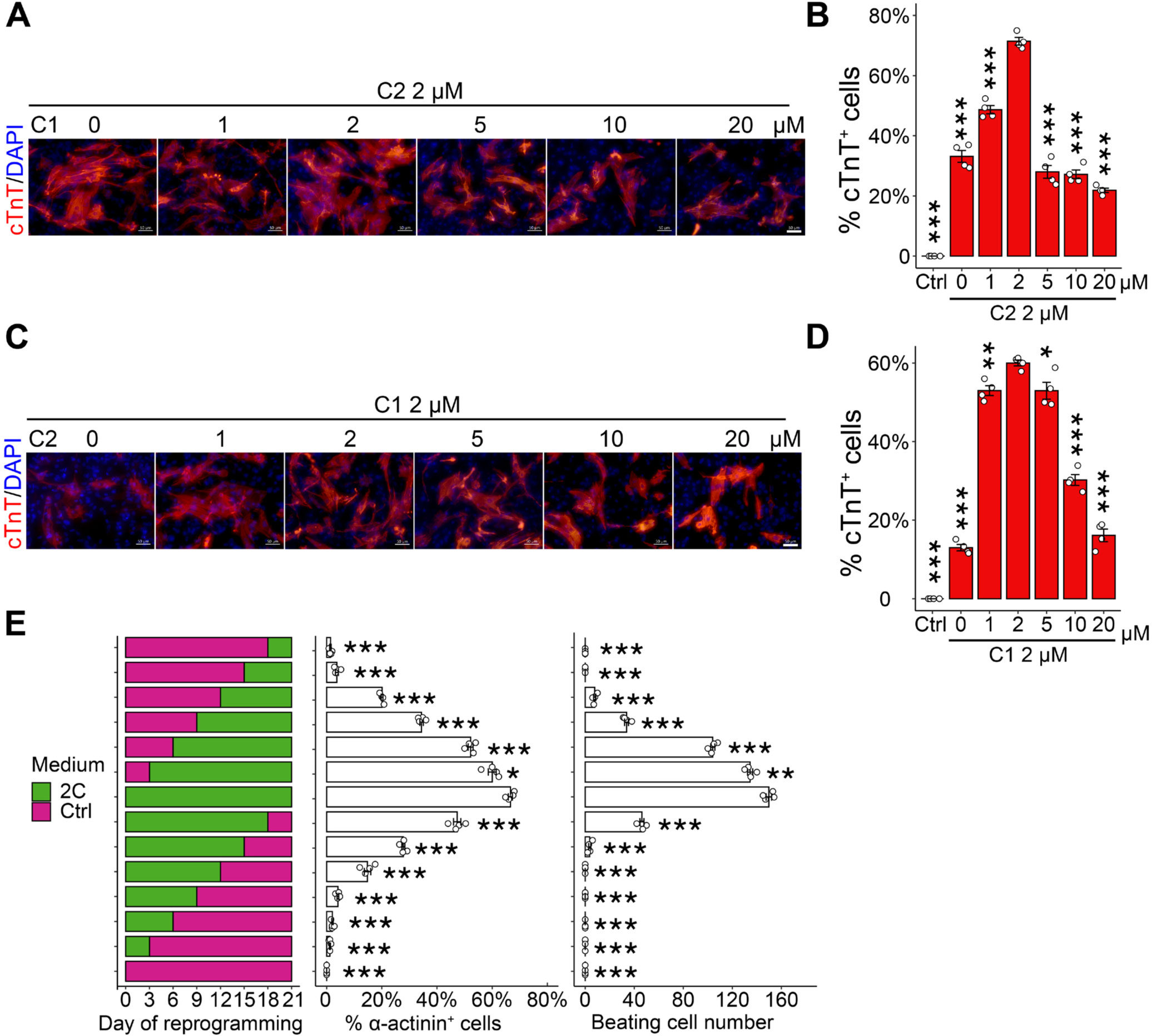
Dosage and timing optimization for 2C, related to Figure 1

**Figure S2.**
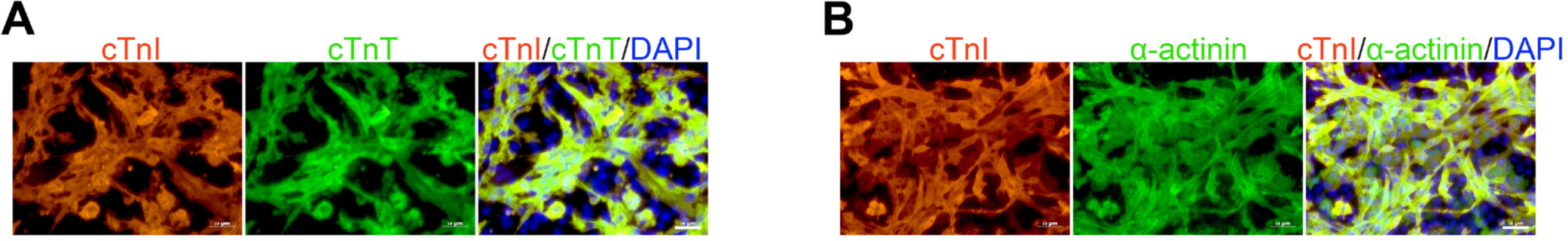
MT+2C induced iCMs co-express multiple cardiomyocyte-specific markers, related to Figure 1

**Figure S3.**
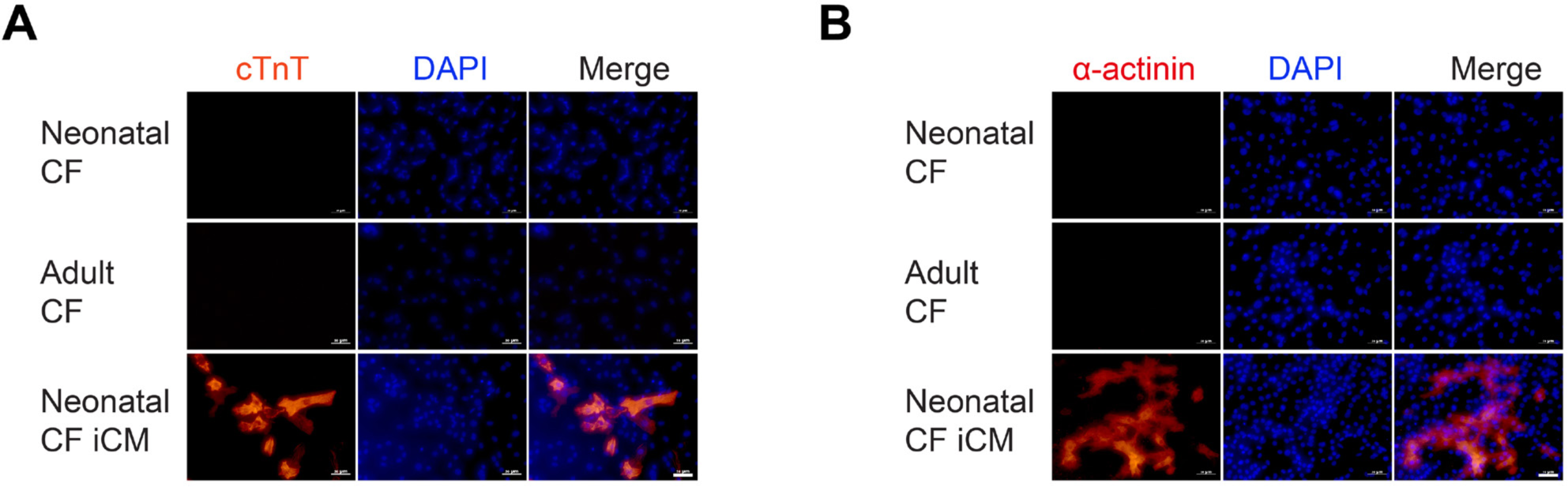
Confirmation of mouse cardiac fibroblasts, related to Figure 2

**Figure S4.**
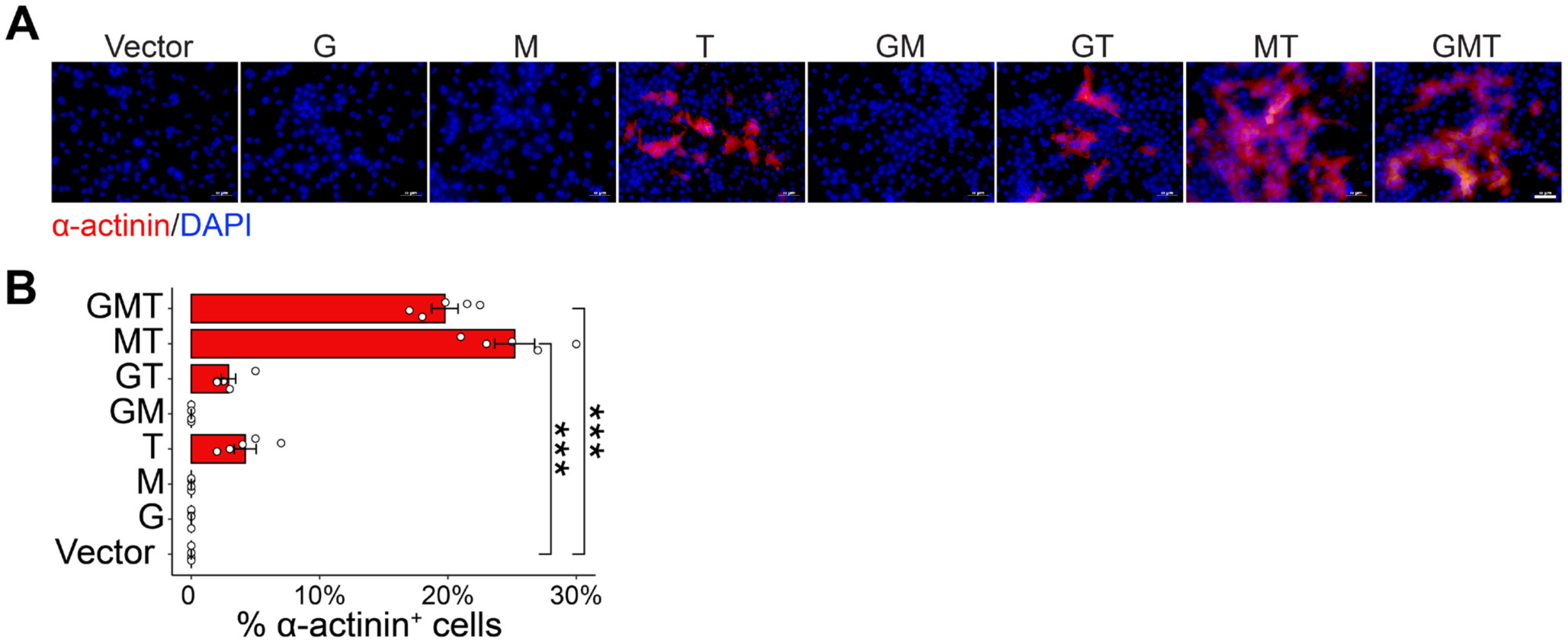
Different combinations of transcription factors reveal MT enables cardiac reprogramming in presence of 2C, related to Figure 2

**Figure S5.**
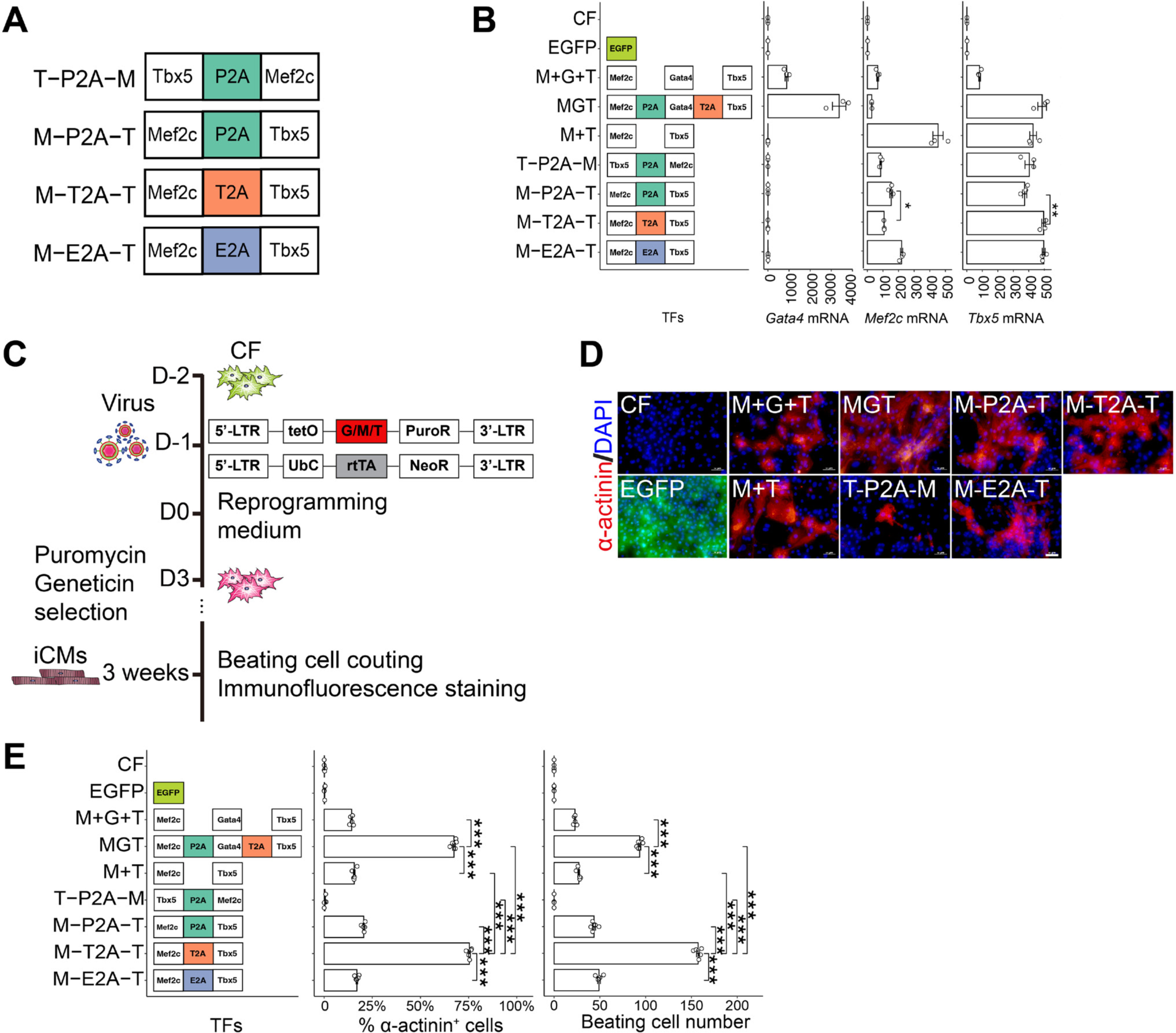
Optimized stoichiometry of Mef2c and Tbx5 results in higher efficiency, related to Figure 2

**Figure S6.**
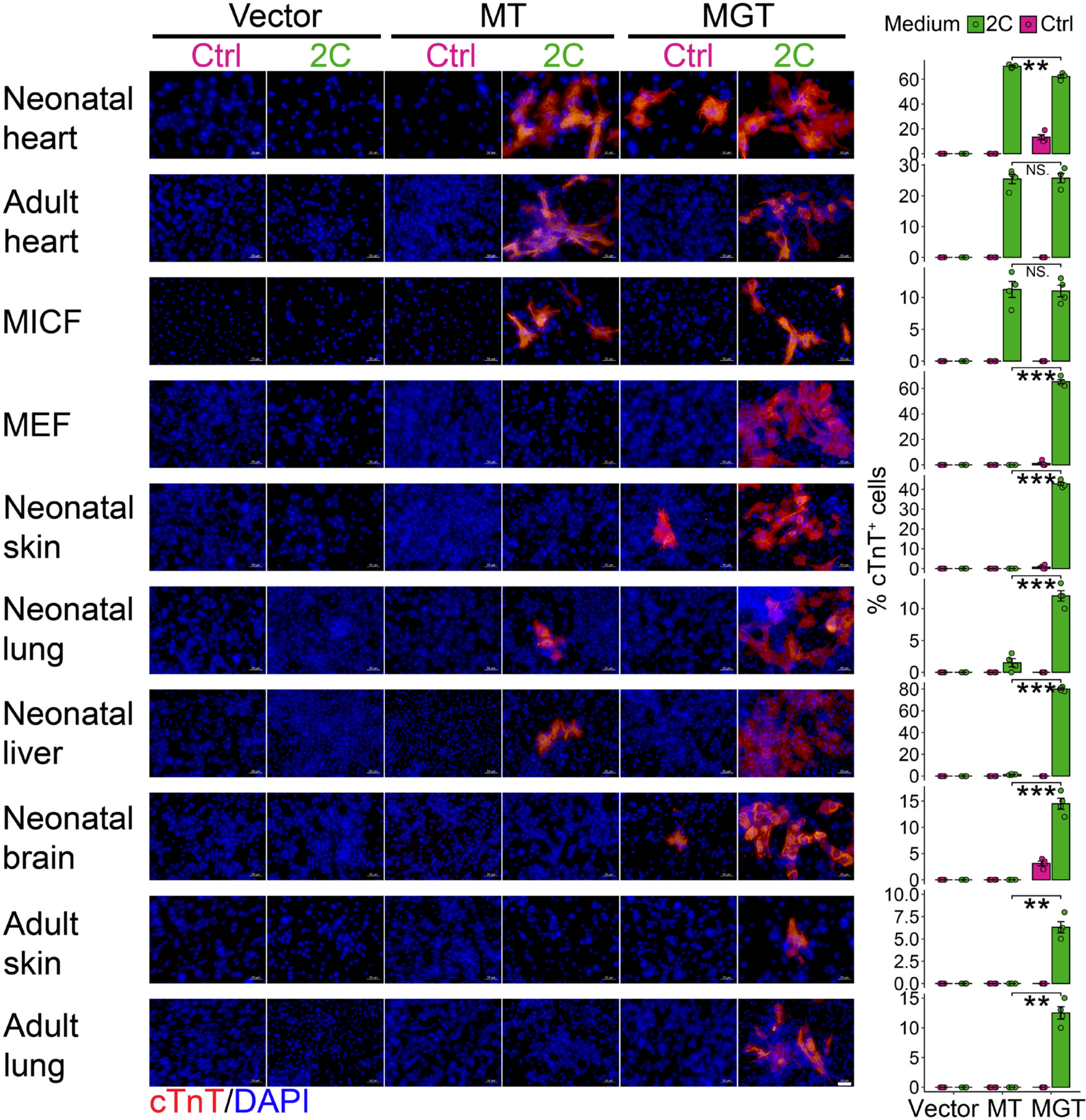
MT+2C selectively reprogram fibroblasts derived heart into iCMs, related to Figure 2

**Figure S7.**
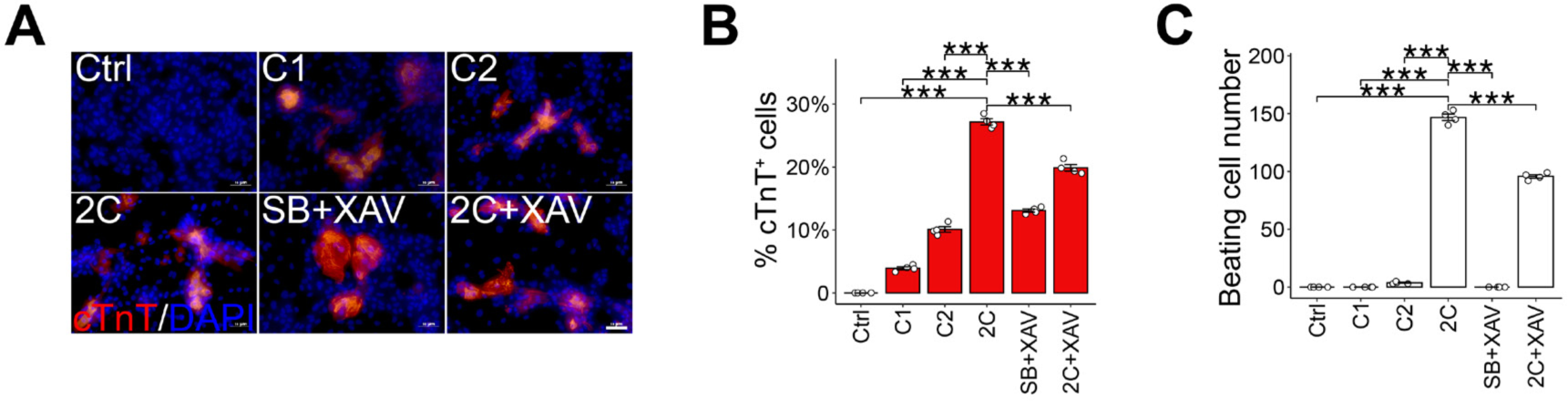
Both C1 and C2 are essential for substituting Gata4, related to Figure 2

**Figure S8.**
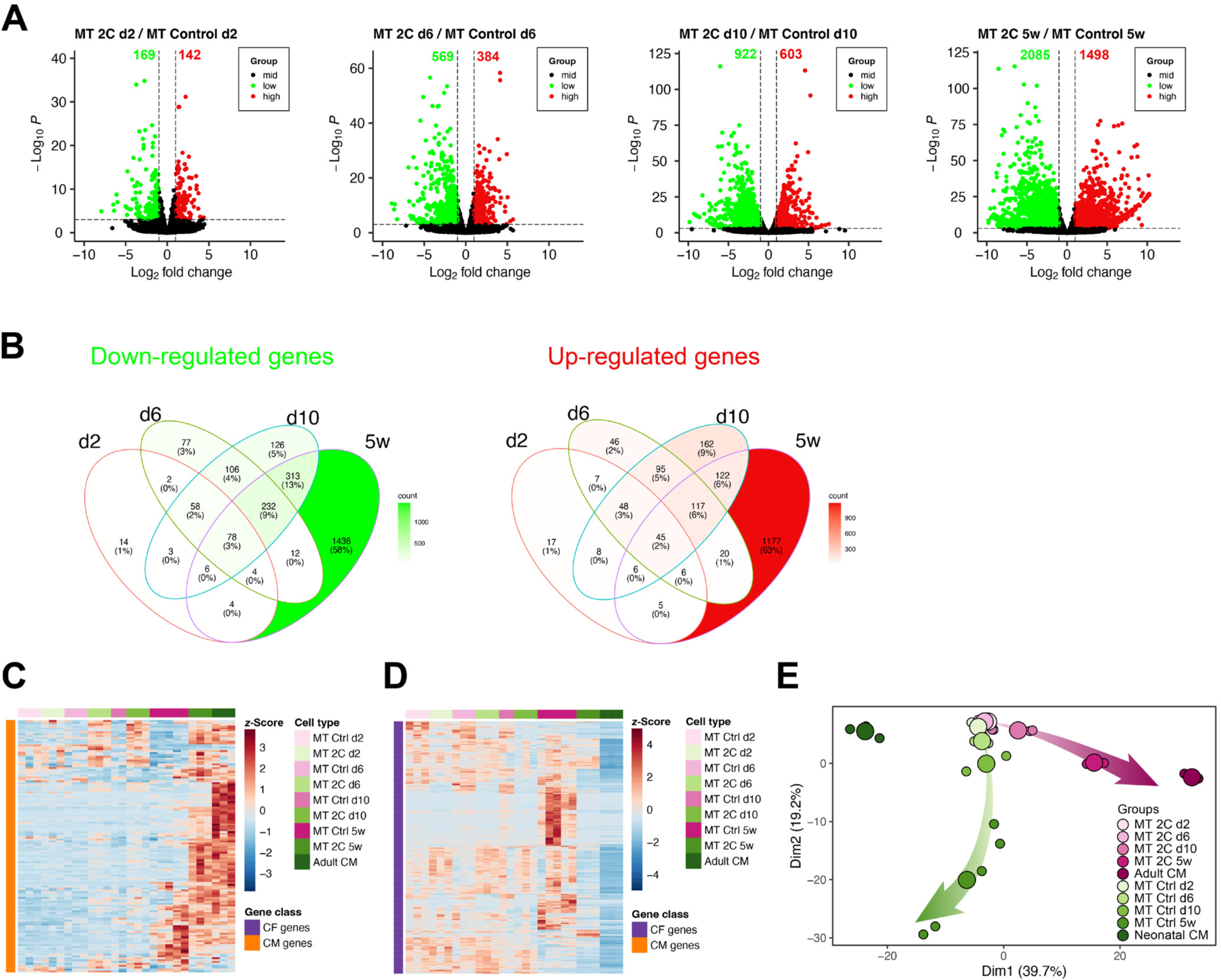
2C enhances cardiac reprogramming progressively, related to Figure 3

**Figure S9.**
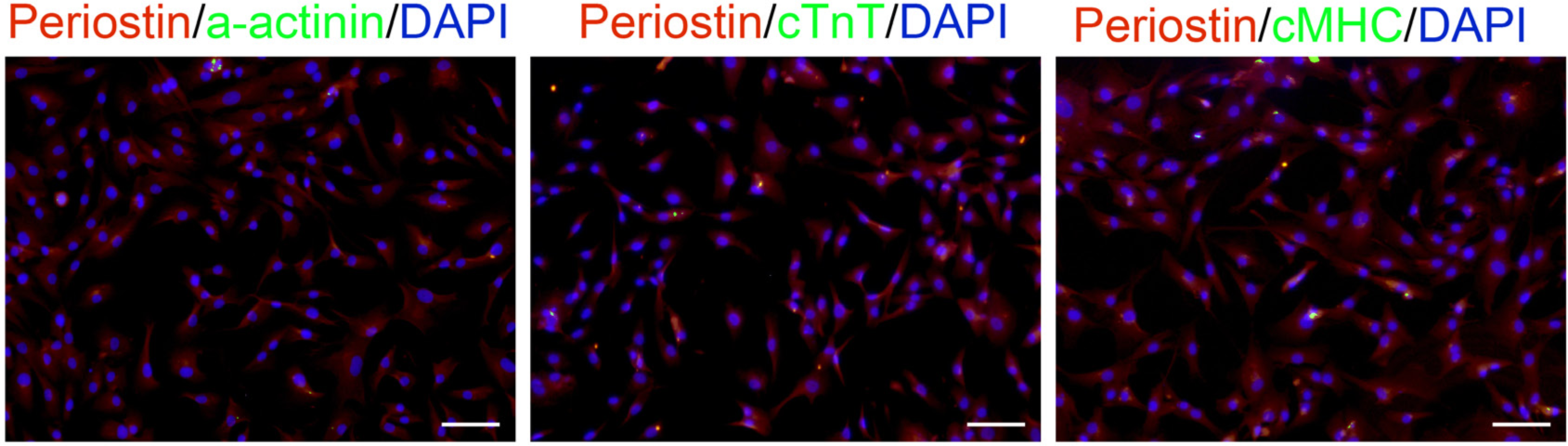
Confirmation of human cardiac fibroblasts, related to Figure 4

**Figure S10.**
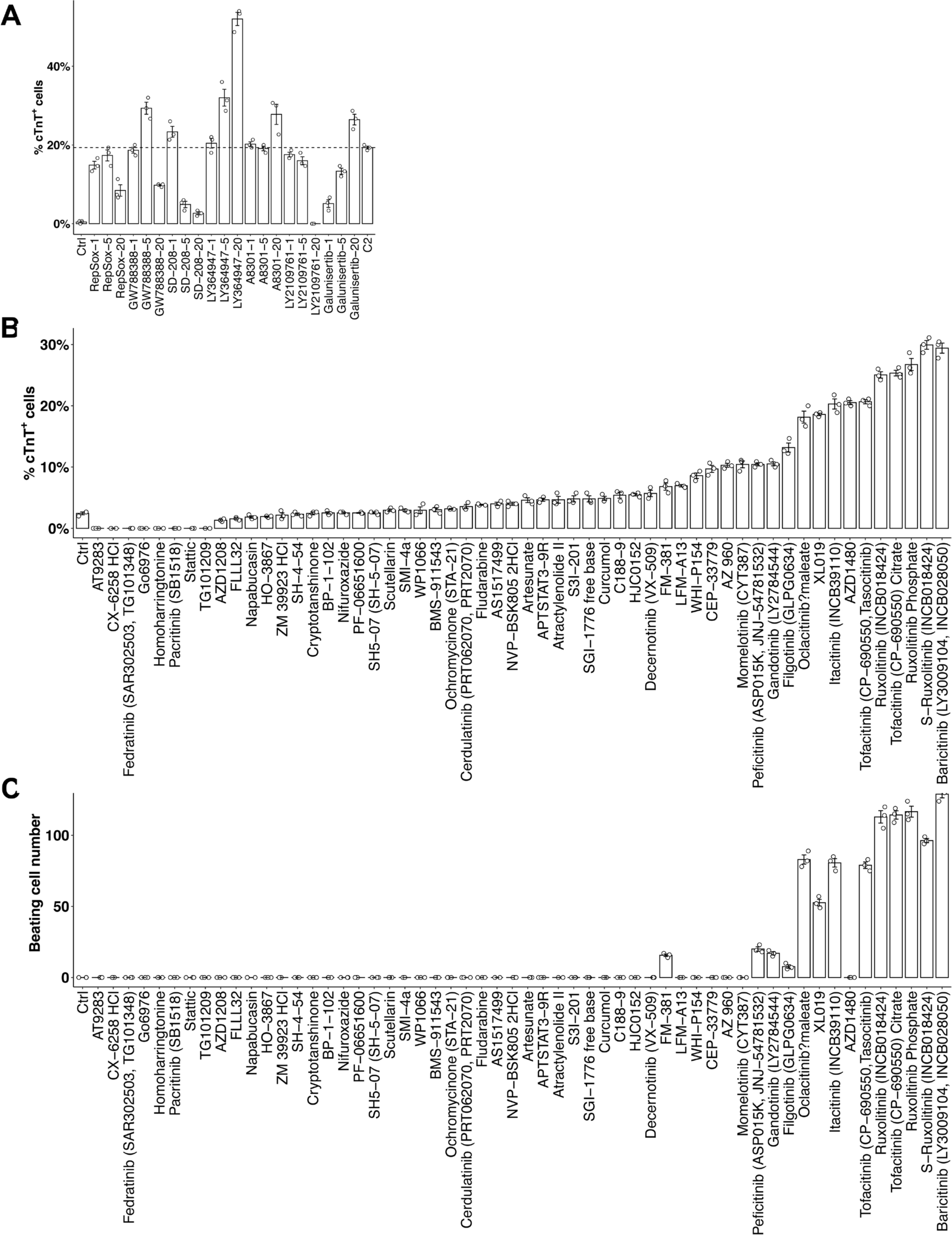
All TGF beta inhibitors tested works as C1, whereas only a few pan-Jak inhitors works as C2, related to Figure 7

**Figure S11.**
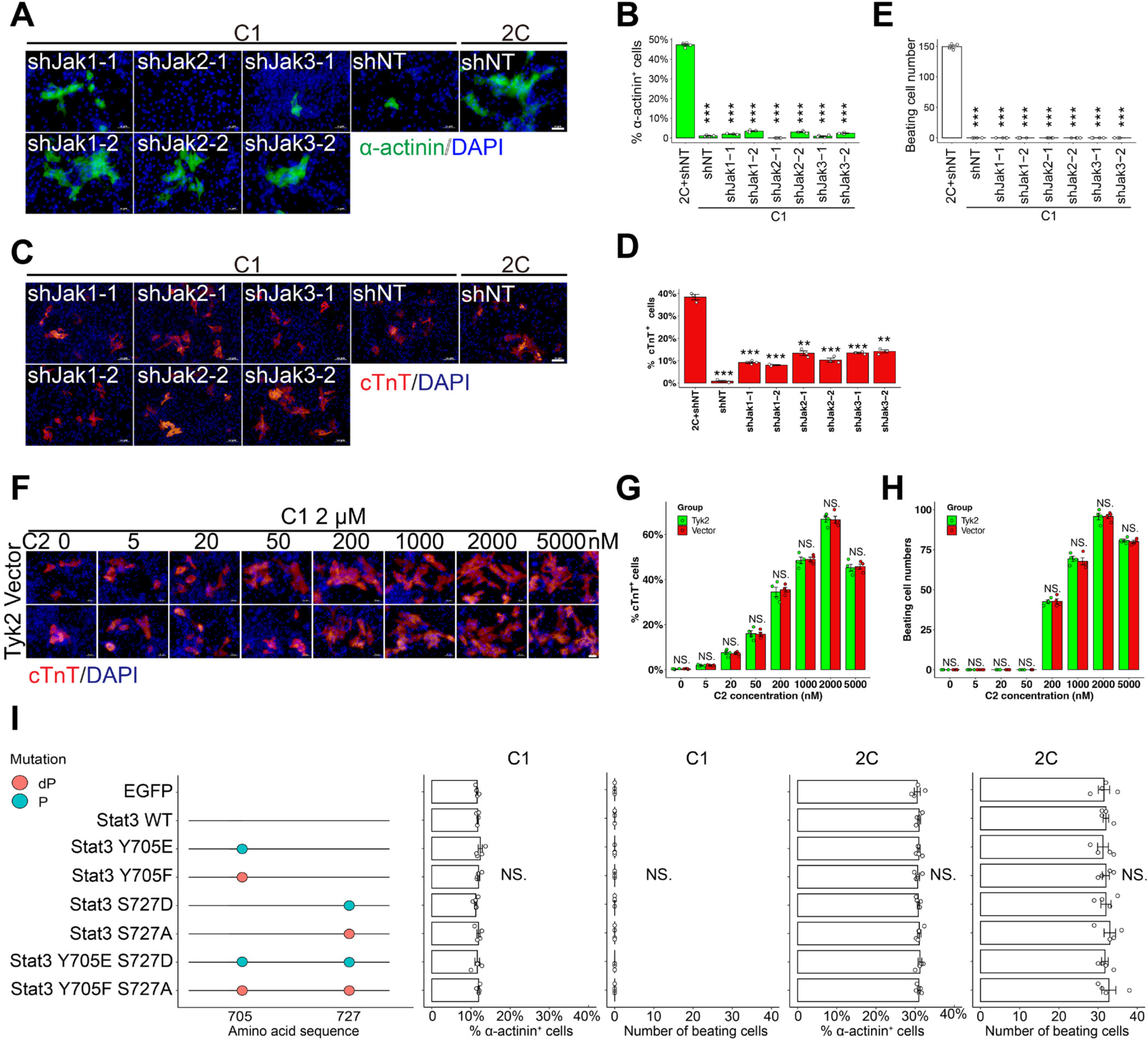
C2 enhances cardiac reprogramming independent of inhibiting the canonical Jak-Stat signal pathway, related to Figure 7

**Figure S12.**
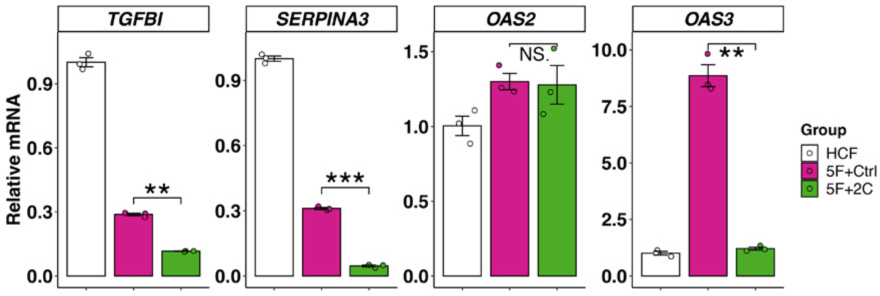
TGFB, SERPINA3, and OAS3 are significantly down-regulated in 2C-treated hiCMs.

