## Supplementary material for "Robust Small Molecule-Aided Cardiac Reprogramming Systems Selective to Cardiac Fibroblasts": figure legend

**Figure 1 The 2C combination boosts cardiac reprogramming**

(A) Schematic representation of the screening strategy for chemicals that enhance cardiac reprogramming.

(B-C) Representative images (B) and quantification (C) for ɑMHc-mCherry+ cells 2 weeks after transduction with GMT (Gata4, Mef2c and Tbx5), in combination with Ctrl (DMSO), C1 (SB431542), C2 (Baricitinib), or 2C (SB431542+Baricitinib) treatment of neonatal mouse skin fibroblasts (NSF). n = 5 independent experiments. Scale bars, 100 µm.

(D) Schematic representation of iCMs reprogramming in wild type mouse skin fibroblasts.

(E-F) Representative immunocytochemistry images (E) and quantification (F) of ɑ-actinin+ cells from NSF 10 days after transduction with GMT, in combination with Ctrl (DMSO), C1, C2, 2C, SB+XAV (SB431542+XAV939), or 2C+XAV (SB431542+Baricitini+XAV) treatment. n = 3 independent experiments.

(G-H) Representative immunocytochemistry images (G) and quantification (H) of cTnT+ cells from NSF 10 days after transduction with GMT, in combination with Ctrl (DMSO), C1, C2, 2C, SB+XAV (SB431542+XAV939), or 2C+XAV (SB431542+Baricitini+XAV) treatment. n = 3 independent experiments.

(I) Representative immunocytochemistry images of Myl2v+ and ɑ-actinin+ cells 2 weeks after transduction with GMT, in combination with 2C treatment of NSF. Scale bars, 50 µm.

(J) Quantification of the number of spontaneous beating cells 2 weeks after transduction with GMT, in combination with Ctrl, 2C, or SB+XAV treatment in NSF. n = 5 independent experiments. All data are presented as the means ± SEM. *p < 0.05, **p < 0.01, ***p < 0.001 versus the relevant control. NS., not significant.

**Figure 2. 2C+MT selectively reprograms cardiac fibroblasts**

(A) Cardiogenic genes expression in adult mouse cardiac fibroblasts (ACF) and adult mouse skin fibroblasts (ASF). n = 2 independent experiments.

(B-D) Representative immunocytochemistry images (B) and quantification of ɑ-actinin+ cells (C), cTnT+ cells (D) 10 days after transduction with vector, MT, or GMT, in combination with Ctrl (DMSO) or 2C treatment of neonatal mouse cardiac fibroblasts. n = 3 independent experiments.

(E) Quantification of the number of spontaneous beating cardiomyocytes over time for groups transduced with vector control, MT, or MGT in various combinations with Ctrl, 2C, SB+XAV, or 2C+XAV treatment. n = 5 independent experiments.

(F) Representative immunocytochemistry images and quantification of cTnT+ cells 3 weeks post transduction with MT or MGT, in combinations with Ctrl or 2C treatment of various fibroblasts derived from indicated organs/tissues. n = 5 independent experiments. Scale bars, 50 µm.

(G) Schematic representation of CD90.2+ cardiac fibroblasts and CD31+ endothelial cells sorting by MACS (magnetic-activated cell sorting) and reprogramming.

(H-I) Representative immunocytochemistry images (H) and quantification (I) of cTnT+ cells 3 weeks post transduction with MT or MGT, in combinations with Ctrl or 2C treatment in CD90.2+ fibroblasts or CD31+ endothelial cells. n = 4 independent experiments. Scale bars, 50 µm.

(J) Schematic representation of cardiac reprogramming and in ACF and ASF.

(K) Heatmap of the relative expression of a set of cardiomyocyte-specific genes in vector+Ctrl, MT+Ctrl, or MT+2C treated cells derived from ACF and ASF, 2 weeks post transduction. n = 2 independent experiments.

(L-N) Representative immunocytochemistry images (L) and quantification (M) of cTnI+ cells and the number of spontaneous beating cells (N) 3 weeks after transduction with MT and shNT (Non-target control) or shGata4 in neonatal cardiac fibroblasts. All groups were treated with 2C. n = 4 independent experiments. Scale bars, 50 µm.

(O) Schematic representation of cardiac reprogramming in neonatal mouse cardiac fibroblasts treated with shNT or shGata4.

(P) Relative mRNA expression level of Gata4, Tnnt2, Myh6, and Actc1 in neonatal cardiac fibroblasts treated with shNT or shGata4 determined by qPCR. n = 3 independent experiments. All data are presented as the means ± SEM. *p < 0.05, **p < 0.01, ***p < 0.001 versus the relevant control. NS., not significant.

**Figure 3. 2C+MT-reprogrammed iCMs Functionally and Transcriptomically Resemble Adult Cardiomyocytes**

(A) Representative immunocytochemistry images of 2C+MT-induced iCMs cultured for 5 weeks display well-organized sarcomere structure. Scale bars, 25 µm.

(B) 2C+MT and 2C+GMT-induced iCMs cultured for 5 weeks show spontaneous Ca^2+^ oscillation. n = 3-5 independent experiments.

(C) 2C+MT- or 2C+GMT-induced iCMs cultured for 5 weeks display action potentials resembling those of adult mouse ventricular cardiomyocytes. n = 3-5 independent experiments.

(D) Volcano map of differentially expressed genes (DEGs) between 2C and control (Ctrl) treatment on MT-transduced cardiac fibroblasts 5 weeks post induction. n = 2-5 independent experiments.

(E) Relative expression of a set of cardiomyocytes (CM) and cardiac fibroblast-related (CF) genes determined by RNA-seq. n = 2-5 independent experiments.

(F) Relative mRNA level of cardiomyocyte-specific gene expression in cells transduced with vector control, MT, or GMT in combination with Ctrl (DMSO) or 2C for 5 weeks, determined by qPCR. ACM represents adult mouse ventricular cardiomyocyte. n = 3 independent experiments.

(G) Pearson correlation analysis of DEGs between 2C and control (Ctrl) treatment on 5-week cultured MT-transduced cardiac fibroblasts. n = 2-5 independent experiments.

(H) Principal component analysis (PCA) of DEGs between 2C and control (Ctrl) treatment on MT-transduced cardiac fibroblasts show that 2C+MT- or 2C+GMT-induced iCMs resemble adult cardiomyocyte. n = 2-5 independent experiments.

(I) Enriched gene ontology (GO) terms for differentially regulated genes in iCMs treated with 2C+MT and Ctrl+MT for 5 weeks. All data are presented as the means ± SEM. *p < 0.05, **p < 0.01, ***p < 0.001 versus the relevant control. NS., not significant.

**Figure 4. 2C enhance cardiac reprogramming in human cardiac fibroblasts**

(A-C) Representative immunocytochemistry images (A), quantification of ɑ-actinin+ cells (B), cTnT+ cells (C) 2 weeks after 5F (GATA4, MEF2C, TBX5, MESP1 and MYOCD) transfection, in combination with Ctrl (DMSO) or 2C treatment of human cardiac fibroblasts. n = 3 independent experiments. Scale bars, 100 µm.

(D) Relative mRNA level of cardiomyocyte-specific genes (MYH6, TNNT2, ACTN2 and ACTC1) determined by qPCR, related to A and B. n = 3 independent experiments.

(E-G) Representative immunocytochemistry images (E), quantification of ɑ-actinin+ cells (F), cTnT+ cells (G) 2 weeks after 5F or 4F (MEF2C, TBX5, MESP1 and MYOCD) transfection, in combination with Ctrl (DMSO) or 2C treatment of human cardiac fibroblasts. n = 3 independent experiments. Scale bars, 100 µm.

(H-J) Representative immunocytochemistry images (H), quantification of ɑ-actinin+ cells (I), cTnT+ cells (J) 2 weeks after 5F or 4F (MEF2C, TBX5, MESP1 and MYOCD) transfection, in combination with Ctrl (DMSO) or 2C treatment of human skin fibroblasts. n = 3 independent experiments. Scale bars, 100 µm.s

All data are presented as the means ± SEM. *p < 0.05, **p < 0.01, ***p < 0.001 versus the relevant control. NS., not significant.

**Figure 5. 2C Increases the Proportion of iCMs and inhibits cell proliferation**

(A) Schematic representation of three models of the enhancement of 2C on cardiac reprogramming and cell proliferation.

(B) Schematic of reprogramming system and detection of cell proliferation.

(C) Representative immunocytochemistry images for Cyclin D1, ɑ-actinin and DAPI at the indicated time points. Scale bars, 50 µm.

(D) Quantification of the absolute number of ɑ-actinin+ cells over time for groups treated with Ctrl (DMSO) or 2C. n = 4 independent experiments.

(E) Quantification of the percentage of Cyclin D1+ cells over time for groups treated with Ctrl (DMSO) or 2C. n = 4 independent experiments.

(F) Quantification of the number of cells (indicated by DAPI+ cells) over time for groups treated with Ctrl (DMSO) or 2C. n = 4 independent experiments.

(G) Quantification of the percentage of ɑ-actinin+ cells over time for groups treated with Ctrl (DMSO) or 2C. n = 4 independent experiments.

(H) Quantification of the number of Cyclin D1+ a-actinin+ cells over time for groups treated with Ctrl (DMSO) or 2C. n = 4 independent experiments.

(I) Schematic of EdU incorporation and Ki67 staining assay.

(J-K) Representative immunocytochemistry images (J) and quantification (K) of cTnI-EdU+ and cTnI+EdU+ cells. n = 5 independent experiments. Scale bars, 20 µm.

(L-M) Representative immunocytochemistry images (L) and quantification (M) of cTnI-Ki67+ and cTnI+Ki67+ cells. n = 5 independent experiments. Scale bars, 20 µm. All data are presented as the means ± SEM. *p < 0.05, **p < 0.01, ***p < 0.001 versus the relevant control. NS., not significant.

**Figure 6. 2C+MT induced iCMs maintain a stable cardiomyocyte fate**

(A) Schematic of reprogramming system to test the stability of MT+2C-induced iCMs.

(B) Representative immunocytochemistry images for cTnI and DAPI at the indicated time points. Scale bars, 50 µm.

(C) Quantification of the number of cTnI+ cells over time for groups treated with Ctrl (DMSO) or 2C. n = 4 independent experiments.

(D) Quantification of the number of cells (indicated by DAPI+ cells) over time for groups treated with Ctrl (DMSO) or 2C. n = 4 independent experiments.

(E) Quantification of the percentage of cTnI+ cells over time for groups treated with Ctrl (DMSO) or 2C. n = 4 independent experiments. All data are presented as the means ± SEM. *p < 0.05, **p < 0.01, ***p < 0.001 versus the relevant control. NS., not significant.

**Figure 7. C2 synergizes cardiomyocyte induction via suppression of C1-activated molecular barriers**

(A-B) Representative immunocytochemistry images (A) and quantification (B) for ɑ-actinin+ cells 3 weeks after MT transduction, in combination with shNT, shAlk4, or shAlk5 and cultured in C2 or 2C medium. n = 4 independent experiments. Scale bars, 50 µm.

(C) Quantification of spontaneous beating cells over time, related to A and B.

(D-F) Representative immunocytochemistry images (D) , quantification of cTnI+ cells (E), and quantification of beating cells 3 weeks after MT transduction, in combination with 2C medium and over-expression of vector (vector control), Alk5 or Tgfb1. n = 4 independent experiments. Scale bars, 50 µm.

(G-H) Representative immunocytochemistry images (G) and quantification of cTnT+ cells (H) 3 weeks after MT transduction, in combination with C1 or 2C medium and knock-down of NT (non-target control) or Tyk2. n = 4 independent experiments. Scale bars, 50 µm.

(I-J) Representative immunocytochemistry images (I) and quantification of ɑ-actinin+ cells (J) 3 weeks after MT transduction, in combination with C1 or 2C medium and knock-down of NT (non-target control) or Tyk2. n = 4 independent experiments. Scale bars, 50 µm.

(K) Quantification of spontaneous beating cells over time, related to G-J. n = 4 independent experiments.

(L) Relative expression of Oas2, Oas3, Serpina3n, and Tgfbi in cardiac fibroblasts transduced with MT, in combination with Ctrl (DMSO) or 2C medium over time. n = 2-3 independent experiments.

(M) Relative expression of Oas2, Oas3, Serpina3n, and Tgfbi in cardiac fibroblasts 3 weeks after transduction with MT, in combination with Ctrl (DMSO), C1 or 2C medium. n = 2-3 independent experiments.

(N-P) Representative immunocytochemistry images (N) and quantification (O) of cTnI+ cells and spontaneous beating cells (P) 3 weeks after transduction with MT and knock-down of NT (Non-target control), 5B (Tyk2 (B1), Oas2 (B2), Oas3 (B3), Serpina3n (B4) and Tgfbi (B5)) and each combination of 4 factors. n = 4 independent experiments. All groups were cultured in C1 medium. Scale bars, 50 µm.

(Q-S) Representative immunocytochemistry images (Q) and quantification (R) of cTnI+ cells and spontaneous beating cells (S) 3 weeks after transduction with MT and over-expression of vector, 5B, and each combination of 4 factors. n = 4 independent experiments. All groups were cultured in 2C medium. Scale bars, 50 µm.

(T) Schematic of the downstream of C1 and C2. All data are presented as the means ± SEM. *p < 0.05, **p < 0.01, ***p < 0.001 versus the relevant control. NS., not significant.

**Supplemental figures legend**

**Figure S1. Dosage and timing optimization for 2C, related to Figure 1**

(A-B) Representative immunocytochemistry images (A) and quantification (B) of cTnT+ cells 2 weeks after transduction with GMT and supplemented with C1 at different concentration. C2 was supplemented at a concentration of 2 µM. n = 4 independent experiments. Scale bars, 50 µm.

(C-D) Representative immunocytochemistry images (C) and quantification (D) of cTnT+ cells 2 weeks after transduction with GMT and supplemented with C2 at different concentrations. C1 was supplemented at a concentration of 2 µM. n = 4 independent experiments. Scale bars, 50 µm.

(E) Quantification of the percentage of α-actinin+ cells and the number of beating cells with different duration of 2C treatment. n = 4 independent experiments. All data are presented as the means ± SEM. *p < 0.05, **p < 0.01, ***p < 0.001 versus the relevant control. NS., not significant.

**Figure S2. MT+2C induced iCMs co-express multiple cardiomyocyte-specific markers, related to Figure 1**

(A) MT+2C induced iCMs co-expressed cTnI and cTnT. Scale bars, 50 µm.

(B) MT+2C induced iCMs co-expressed cTnI and α-actinin. Scale bars, 50 µm.

**Figure S3 Confirmation of mouse cardiac fibroblasts, related to Figure 2**

(A-B) Neonatal mouse cardiac fibroblasts and adult mouse cardiac fibroblasts were validated with cTnT (A) and ⍺-actinin (B) to exclude any contamination with cardiomyocytes.

**Figure S4. Different combinations of transcription factors reveal MT enables cardiac reprogramming in presence of 2C, related to Figure 2**

(A) Representative immunocytochemistry images of α-actinin+ cells 3 weeks after transduction with different combinations of transcription factors, supplemented with 2C medium. Scale bars, 50 µm.

(B) Quantification of α-actinin+ cells 3 weeks after transduction with different combinations of transcription factors, supplemented with 2C medium. n = 5 independent experiments. All data are presented as the means ± SEM. *p < 0.05, **p < 0.01, ***p < 0.001 versus the relevant control. NS., not significant.

**Figure S5 Optimized stoichiometry of Mef2c and Tbx5 results in higher efficiency, related to Figure 2**

(A) Schematic of bi-cistronic constructs with different 2A sequences.

(B) Relative expression of Mef2c, Tbx5, and Gata4 in different bi-cistronic constructs, determined by qPCR. n = 3 independent experiments.

(C) Workflow of stoichiometry optimization of Mef2c and Tbx5.

(D-E) Representative immunocytochemistry images (D) and quantification (E) of α-actinin+ cells and the number of spontaneous beating cells (E) 3 weeks after transduction with the indicated construct in cardiac fibroblasts. n = 5 independent experiments. Scale bars, 50 µm. All data are presented as the means ± SEM. *p < 0.05, **p < 0.01, ***p < 0.001 versus the relevant control. NS., not significant.

**Figure S6 MT+2C selectively reprogram fibroblasts derived from heart into iCMs, related to Figure 2**

Representative immunocytochemistry images and quantification of cTnT+ cells 3 weeks post transduction with MT or MGT, in combinations with Ctrl or 2C treatment of various fibroblasts derived from indicated organs/tissues. n = 5 independent experiments. Scale bars, 50 µm. MICF, adult mouse cardiac fibroblasts isolated from mice treated with myocardial infarction. MEF, mouse embryonic fibroblasts.

**Figure S7. Both C1 and C2 are essential for substituting Gata4, related to Figure 2**

(A-C) Representative immunocytochemistry images (A), quantification of cTnT+ cells (B) and quantification of the number of spontaneous beating cells (C) 3 weeks after transduction with MT, in combination with Ctrl (DMSO), C1, C2, 2C, SB+XAV (SB43542+XAV939), or 2C+XAV (SB431542+Baricitinib+XAV939). Scale bars, 50 µm. n = 4 independent experiments. All data are presented as the means ± SEM. *p < 0.05, **p < 0.01, ***p < 0.001 versus the relevant control. NS., not significant.

**Figure S8. 2C enhances cardiac reprogramming progressively, related to Figure 3**

(A) Volcano map of differentially expressed genes (DEGs) between MT+2C and MT+Ctrl treatment of cardiac fibroblasts on d2, d6, d10, and 5 weeks after induction. n = 2-3 independent experiments.

(B) Venn diagram of down-regulated genes and up-regulated genes in (A).

(C) Relative expression of cardiomyocyte (CM) related genes in cultured cells with MT+Ctrl and MT+2C treatment d2, d6, d10, and 5 weeks after induction. n = 2-3 independent experiments.

(D) Relative expression of a set of cardiac fibroblast (CF) related genes in cultured cells with MT+Ctrl and MT+2C treatment d2, d6, d10, and 5 weeks after induction. n = 2-3 independent experiments.

(E) Principal component analysis (PCA) of DEGs between 2C and control (Ctrl) treatment of MT-transduced cardiac fibroblasts on d2, d6, d10, and 5 weeks after induction. n = 2-3 independent experiments.

**Figure S9. Confirmation of human cardiac fibroblasts, related to Figure 4**

Human cardiac fibroblasts (HCF) were confirmed by the expression of Periostin and without expression of a-actinin, cTnT or cMHC. Scale bars, 100 µm.

**Figure S10. All TGF beta inhibitors tested works as C1, whereas only a few pan-Jak inhibitors works as C2, related to Figure 7**

(A) Quantification for cTnT+ cells 3 weeks after transduction with MT, in combination with C2 and indicated TGF beta inhibitors at different concentrations. n = 3 independent experiments.

(B) Quantification for cTnT+ cells 3 weeks after transduction with MT, in combination with C1 and indicated Jak inhibitors. n = 3 independent experiments.

(C) Quantification of the number of spontaneous beating cells 3 weeks after transduction with MT, in combination with C1 and indicated Jak inhibitors. n = 3 independent experiments.

**Figure S11. C2 enhances cardiac reprogramming independent of inhibiting the canonical Jak-Stat signal pathway, related to Figure 7**

(A-B) Representative immunocytochemistry images (A) and quantification of α-actinin+ cells (B) 3 weeks after transduction with MT, supplemented with C1 treatment. n = 4 independent experiments. Scale bars, 50 µm.

(C-D) Representative immunocytochemistry images (C) and quantification of cTnT+ cells (D) 3 weeks after transduction with MT, supplemented with C1 treatment. n = 3 independent experiments. Scale bars, 50 µm.

(E) Quantification of the number of spontaneous beating cells 3 weeks after transduction with MT, supplemented with C1 treatment. n = 4 independent experiments.

(F-H) Representative immunocytochemistry images (F) , quantification of cTnT+ cells (G) and spontaneous beating cells (H) 3 weeks after transduction with MT, supplemented with C1 treatment and C2 at indicated concentrations. n = 4 independent experiments. Scale bars, 50 µm.

(I) Quantification of the percentage of α-actinin+ cells and the number of spontaneous beating cells 3 week after transduction with MT and indicated EGFP or Stat3 mutants, cultured in C1 or 2C medium as indicated in the figure. P and dP represent mutants at indicated amino acids to simulate the phosphorylated and dephosphorylated states of Stat3. n = 4 independent experiments. All data are presented as the means ± SEM. *p < 0.05, **p < 0.01, ***p < 0.001 versus the relevant control. NS., not significant.

**Supplemental movies legend**

Movie S1 2C treated iCMs are cross-linked and difficult to dissociate.

Movie S2 No spontaneous beating cell could be detected in NSF that treated with GMT+Ctrl for 2 weeks, related to Figure 1L.

Movie S3 Spontaneous beating cells detected in NSF that treated with GMT+2C for 2 weeks, related to Figure 1L.

Movie S4 Spontaneous beating cells detected in NSF that treated with GMT+SB431542+XAV939 for 2 weeks, related to Figure 1L.

Movie S5 Spontaneous beating cells detected in neonatal cardiac fibroblasts that treated with MGT+2C for 3 weeks, related to Figure 2D.

Movie S6 Spontaneous beating cells detected in neonatal cardiac fibroblasts that treated with MT+2C for 3 weeks, related to Figure 2D.

Movie S7 Spontaneous beating rod-shaped iCMs detected in neonatal cardiac fibroblasts that treated with MT+2C for 5 weeks, related to Figure 3A.

Movie S8 Calcium transients detected in neonatal cardiac fibroblasts that treated with MT+2C for 5 weeks, related to Figure 3B.
