## Supplementary material for "Robust Small Molecule-Aided Cardiac Reprogramming Systems Selective to Cardiac Fibroblasts": method

**KEY RESOURCES TABLE**

The table highlights the reagents, genetically modified organisms and strains, cell lines, software, instrumentation, and source data **essential** to reproduce results presented in the manuscript. Depending on the nature of the study, this may include standard laboratory materials (i.e., food chow for metabolism studies, support material for catalysis studies), but the table is **not** meant to be a comprehensive list of all materials and resources used (e.g., essential chemicals such as standard solvents, SDS, sucrose, or standard culture media do not need to be listed in the table). **Items in the table must also be reported in the method details section within the context of their use.** To maximize readability, the number of **oligonucleotides and RNA sequences** that may be listed in the table is restricted to no more than 10 each. If there are more than 10 oligonucleotides or RNA sequences to report, please provide this information as a supplementary document and reference the file (e.g., See Table S1 for XX) in the key resources table.

***Please note that ALL references cited in the key resources table must be included in the references list.*** Please report the information as follows:

- **REAGENT or RESOURCE:** Provide full descriptive name of the item so that it can be identified and linked with its description in the manuscript (e.g., provide version number for software, host source for antibody, strain name). In the experimental models section (applicable only to experimental life science studies), please include all models used in the paper and describe each line/strain as: model organism: name used for strain/line in paper: genotype. (i.e., Mouse: OXTR^fl/fl^: B6.129(SJL)-Oxtr^tm1.1Wsy/J^). In the biological samples section (applicable only to experimental life science studies), please list all samples obtained from commercial sources or biological repositories. Please note that software mentioned in the methods details or data and code availability section needs to also be included in the table. See the sample tables at the end of this document for examples of how to report reagents.
- **SOURCE:** Report the company, manufacturer, or individual that provided the item or where the item can be obtained (e.g., stock center or repository). For materials distributed by Addgene, please cite the article describing the plasmid and include “Addgene” as part of the identifier. If an item is from another lab, please include the name of the principal investigator and a citation if it has been previously published. If the material is being reported for the first time in the current paper, please indicate as “this paper.” For software, please provide the company name if it is commercially available or cite the paper in which it has been initially described.
- **IDENTIFIER:** Include catalog numbers (entered in the column as “Cat#” followed by the number, e.g., Cat#3879S). Where available, please include unique entities such as [RRIDs](https://www.force11.org/group/resource-identification-initiative), Model Organism Database numbers, accession numbers, and PDB, CAS, or CCDC IDs. For antibodies, if applicable and available, please also include the lot number or clone identity. For software or data resources, please include the URL where the resource can be downloaded. Please ensure accuracy of the identifiers, as they are essential for generation of hyperlinks to external sources when available. Please see the Elsevier [list of data repositories](https://www.elsevier.com/authors/author-resources/research-data/data-base-linking) with automated bidirectional linking for details. When listing more than one identifier for the same item, use semicolons to separate them (e.g., Cat#3879S; RRID: AB_2255011). If an identifier is not available, please enter “N/A” in the column.
  - ***A NOTE ABOUT RRIDs:*** We highly recommend using RRIDs as the identifier (in particular for antibodies and organisms but also for software tools and databases). For more details on how to obtain or generate an RRID for existing or newly generated resources, please [visit the RII](https://www.force11.org/group/resource-identification-initiative) or [search for RRIDs](https://scicrunch.org/resources).

Please use the empty table that follows to organize the information in the sections defined by the subheading, skipping sections not relevant to your study. Please do not add subheadings. To add a row, place the cursor at the end of the row above where you would like to add the row, just outside the right border of the table. Then press the ENTER key to add the row. Please delete empty rows. Each entry must be on a separate row; do not list multiple items in a single table cell. Please see the sample tables at the end of this document for relevant examples in the life and physical sciences of how reagents and instrumentation should be cited.

***TABLE FOR AUTHOR TO COMPLETE***

*Please upload the completed table as a separate document.* ***Please do not add subheadings to the key resources table.*** *If you wish to make an entry that does not fall into one of the subheadings below, please contact your handling editor.* ***Any subheadings not relevant to your study can be skipped.*** *(****NOTE:*** *For authors publishing in Cell Genomics, Cell Reports Medicine, Current Biology, and Med, please note that references within the KRT should be in numbered style rather than Harvard.)*

**Key resources table**

| REAGENT or RESOURCE | SOURCE | IDENTIFIER |
| --- | --- | --- |
| Antibodies | | |
| Cardiac Troponin T Monoclonal Antibody (13-11) | Thermo Fisher | Cat#MA5-12960 |
| Anti-Cardiac Troponin I | abcam | Cat#ab56357 |
| Anti-Sarcomeric Alpha Actinin | abcam | Cat#ab9465 |
| Anti-Cyclin D1 | abcam | Cat#ab16663 |
| Anti-Myosin Light Chain 2 | abcam | Cat#ab48003 |
| Donkey anti-Mouse Alexa Fluor 488 | Thermo Fisher | Cat#A-21202 |
| Donkey anti-Mouse Alexa Fluor 555 | Thermo Fisher | Cat#A-31570 |
| Donkey anti-Rabbit Alexa Fluor 488 | Thermo Fisher | Cat#A-21206 |
| Donkey anti-Rabbit Alexa Fluor 555 | Thermo Fisher | Cat#A-31572 |
| Donkey anti-Goat Alexa Fluor 488 | Thermo Fisher | Cat#A-11055 |
| Donkey anti-Goat Alexa Fluor 555 | Thermo Fisher | Cat#A-21432 |
| CD90.2 MicroBeads | Miltenyi | Cat#130-121-278 |
| Bacterial and virus strains | | |
| Biological samples |  |  |
| Chemicals, peptides, and recombinant proteins | | |
| SB431542 | Selleck | Cat#S1067 |
| Baricitinib | Selleck | Cat#S2851 |
| Doxycycline hyclate | Sigma-Aldrich | Cat#D9891 |
| XAV939 | Selleck | Cat#S1180 |
| Critical commercial assays | | |
| pEASY®-Uni Seamless Cloning and Assembly Kit | TransGen | Cat#CU101 |
| RNeasy Plus Mini Kit | QIAGEN | Cat#74134 |
| QIAGEN Plasmid Mega Kit | QIAGEN | Cat#12183 |
| Fluo4-AM | Thermo Fisher | Cat#F14201 |
| TransScript® One-Step gDNA Removal and cDNA Synthesis SuperMix | TransGen | Cat#AT311 |
| ChamQ SYBR qPCR Master Mix(Without ROX) | Vazyme | Cat# Q321-02 |
| Deposited data | | |
| RNA-seq | GEO Database | GSE213021 |
| Experimental models: Cell lines | | |
| NHCF | Lonza | Cat#CC2904 |
| BJ | ATCC | Cat#CRL-2522 |
| Experimental models: Organisms/strains | | |
| Mouse: B6;D2-Tg(Myh6*-mCherry)2Mik/J | The Jackson Laboratory | Mouse strain #: 021577 |
| Mouse: CD-1(ICR) | Beijing Vital River LaboratoryAnimal Technology Co., Ltd | Mouse strain #: 201 |
| Oligonucleotides | | |
| Separated attachement, Table S1. |  |  |
| Recombinant DNA | | |
| FU-tet-o-Puro-Vector | This paper | N/A |
| FU-tet-o-Puro-EGFP | This paper | N/A |
| FU-tet-o-Puro-Gata4 | This paper | N/A |
| FU-tet-o-Puro-Mef2c | This paper | N/A |
| FU-tet-o-Puro-Tbx5 | This paper | N/A |
| FU-tet-o-Puro-MGT | This paper | N/A |
| FU-tet-o-Puro-T-P2A-M | This paper | N/A |
| FU-tet-o-Puro-M-P2A-T | This paper | N/A |
| FU-tet-o-Puro-M-T2A-T | This paper | N/A |
| FU-tet-o-Puro-M-E2A-T | This paper | N/A |
| tetO-hGATA4 | addgene | Cat#46030 |
| tetO-hMEF2C | addgene | Cat#46031 |
| tetO-hTBX5 | addgene | Cat#46032 |
| FU-tet-o-hMESP1 | This paper | N/A |
| FU-tet-o-hMYOCD | This paper | N/A |
| FU-Ubc-rtTA-Neo | This paper | N/A |
| pLenti-BleoR-Vector | This paper | N/A |
| pLenti-BleoR-EGFP | This paper | N/A |
| pLenti-BleoR-Stat3 | This paper | N/A |
| pLenti-BleoR-Stat3-S727A | This paper | N/A |
| pLenti-BleoR-Stat3-S727D | This paper | N/A |
| pLenti-BleoR-Stat3-Y705E | This paper | N/A |
| pLenti-BleoR-Stat3-Y705E S727D | This paper | N/A |
| pLenti-BleoR-Stat3-Y705F | This paper | N/A |
| pLenti-BleoR-Stat3-Y705F S727A | This paper | N/A |
| pLKO.1-NT | Sigma-Aldrich | SHC016H |
| pLKO.1-Jak1-1 | Sigma-Aldrich | TRCN0000023290 |
| pLKO.1-Jak1-2 | Sigma-Aldrich | TRCN0000023292 |
| pLKO.1-Jak2-1 | Sigma-Aldrich | TRCN0000023650 |
| pLKO.1-Jak2-2 | Sigma-Aldrich | TRCN0000023652 |
| pLKO.1-Jak3-1 | Sigma-Aldrich | TRCN0000023475 |
| pLKO.1-Jak3-2 | Sigma-Aldrich | TRCN0000023478 |
| pLKO.1-Tyk2-1 | Sigma-Aldrich | TRCN0000025886 |
| pLKO.1-Tyk2-2 | Sigma-Aldrich | TRCN0000025942 |
| pLKO.1-Tyk2-3 | Sigma-Aldrich | TRCN0000025964 |
| pLKO.1-Tyk2-4 | Sigma-Aldrich | TRCN0000025903 |
| FU-tet-o-BSD-Tyk2 | This paper | N/A |
| pLKO.1-Alk4-1 | Sigma-Aldrich | TRCN0000022574 |
| pLKO.1-Alk4-2 | Sigma-Aldrich | TRCN0000022575 |
| pLKO.1-Alk5-1 | Sigma-Aldrich | TRCN0000322043 |
| pLKO.1-Alk5-2 | Sigma-Aldrich | TRCN0000322044 |
| FU-tet-o-BSD-Vector | This paper | N/A |
| FU-tet-o-BSD-Alk5 | This paper | N/A |
| FU-tet-o-BSD-Tgfb1 | This paper | N/A |
| pLKO.1-Oas2-1 | Sigma-Aldrich | TRCN0000075813 |
| pLKO.1-Oas2-2 | Sigma-Aldrich | TRCN0000075814 |
| pLKO.1-Oas3-1 | Sigma-Aldrich | TRCN0000075823 |
| pLKO.1-Oas3-2 | Sigma-Aldrich | TRCN0000075824 |
| pLKO.1-Tgfbi-1 | Sigma-Aldrich | TRCN0000054809 |
| pLKO.1-Tgfbi-2 | Sigma-Aldrich | TRCN0000054810 |
| pLKO.1-Serpina3n-1 | Sigma-Aldrich | TRCN0000080455 |
| pLKO.1-Serpina3n-2 | Sigma-Aldrich | TRCN0000080456 |
| FU-tet-o-BSD-Oas2 | This paper | N/A |
| FU-tet-o-BSD-Oas3 | This paper | N/A |
| FU-tet-o-BSD-Tgfbi | This paper | N/A |
| FU-tet-o-BSD-Serpina3n | This paper | N/A |
| pLKO.1-Gata4-1 | Sigma-Aldrich | TRCN0000095214 |
| pLKO.1-Gata4-2 | Sigma-Aldrich | TRCN0000095215 |
| pLKO.1-Gata4-3 | Sigma-Aldrich | TRCN0000095216 |
| Software and algorithms | | |
| R version 4.2.1 | R Project | https://www.r-project.org |
| RStudio Desktop 2022.07.1+554 | Rstudio | https://www.rstudio.com |
| DESeq2 1.36.0 | Love et al., 2014 | https://bioconductor.org/packages/release/bioc/html/DESeq2.html |
| tidyverse | tidyverse | https://www.tidyverse.org |
| EnhancedVolcano 1.14.0 | EnhancedVolcano | https://github.com/kevinblighe/EnhancedVolcano |
| ImageJ-Fiji | National Institutes of Health, USA | https://fiji.sc/ |
| Other | | |
| DMEM，high glucose | HyClone | Cat#SH30022.01 |
| MEDIUM 199 | Thermo Fisher | Cat#11150-059 |
| KnockOut Serum Replacement (KSR) | Thermo Fisher | Cat#A3181502 |
| Fetal Bovine Serum (FBS) | VISTECH | Cat#SE100-011 |
| GlutaMAX | Thermo Fisher | Cat#35050–061 |
| MEM Non-Essential Amino Acids | Thermo Fisher | Cat#11140-050 |
| Penicillin-streptomycin | Thermo Fisher | Cat#15140-122 |
| DAPI | Sigma-Aldrich | Cat#D9542 |
| Polybrene | Sigma-Aldrich | Cat#H9268-10G |
| PBS | Corning | Cat#21-040-CVR |
| Trypsin-EDTA | Thermo Fisher | Cat#25200-056 |
| Puromycin | InvivoGen | Cat#ant-pr-1 |
| Geneticin | InvivoGen | Cat#ant-gn-1 |
| Blasticidin | InvivoGen | Cat#ant-bl-05 |
